# It’s about time! Leaf minimum conductance determines time to reach critical thresholds for leaf dehydration in a seasonal tropical forest

**DOI:** 10.1101/2025.07.07.663429

**Authors:** Ron Sunny, Malavika Venu, Bandaru Peddiraju, Souparna Chakrabarty, Deepak Barua

## Abstract

- Leaf minimum conductance (*g*_min_) is important in determining plant responses to drought. However, we do not understand how *g*_min_ is related to drought tolerance across species, and how important it is in determining the time to reach critical dehydration levels.
- In 18 coexisting species from a seasonal tropical forest we quantified *g*_min_ to test relationships with early, moderate and severe dehydration thresholds associated with tolerance to turgor loss, breakdown of structural integrity, and disruption of cellular function, respectively. We quantified other functional and hydraulic traits to determine the major axes of trait variation in these species.
- Variation in *g*_min_ was the primary determinant of the time to reach critical levels of dehydration, and was unrelated to thresholds for early and severe dehydration. Surprisingly, *g*_min_ was negatively related to maximum stomatal conductance, but unrelated to other functional and hydraulic traits.
- These results highlight the importance of avoiding dehydration via minimizing *g*_min_, and suggest that avoidance, tolerance to early, and severe dehydration represent independent strategies for coping with drought. This would allow coexisting species to balance opportunities for carbon gain with costs of physiological breakdown when faced with varying intensities and durations of drought.

## Introduction

Leaf water status is central to determining multiple aspects of leaf function. Increased leaf dehydration during drought causes the breakdown of vital physiological processes with important implications for carbon assimilation, growth, and mortality (Choat *et al*., 2018; Powers *et al*., 2020; McDowell *et al*., 2022). Determining the key physiological processes (Blackman *et al*., 2010; Klein *et al*., 2014; Bartlett *et al*., 2016; Trueba *et al*., 2019), the order in which these are disrupted (Bartlett *et al*., 2016; Trueba *et al*., 2019), and how these differ across species has provided a better mechanistic understanding of species responses to drought. However, a comprehensive understanding of the diverse processes that drive responses to drought is still lacking (Mantova *et al*., 2022), and the ability to predict drought-induced reductions in function, and associated changes in growth, productivity, and mortality remains challenging (Trugman *et al*., 2021). The dynamics of leaf water loss during drought, which determine the time taken to reach critical levels of dehydration has been highlighted as an important aspect of species responses for which we lack essential information (Hammond & Adams, 2019; Brodribb *et al*., 2020; Machado *et al*., 2021; Blackman *et al*., 2023). In this study, we examined leaf water loss during dehydration, and quantified leaf minimum conductance in coexisting evergreen and deciduous species from a seasonally dry tropical forest, and tested how residual water loss from leaves during drought is related to tolerance to early, moderate and severe dehydration.

Over the last couple of decades, a large body of work has identified the key physiological processes that are disrupted with increasing dehydration in leaves, and this has allowed a better understanding of the underlying mechanisms, and the functional consequences of exposure to drought. During the early stages of leaf dehydration, loss of turgor and stomatal closure results in the cessation of carbon assimilation (Klein *et al*., 2014; Bartlett *et al*., 2016). With increasing water loss there is increased leaf shrinkage, breakdown in the structural integrity of leaves, and the loss of conductance of the extra-xylem mesophyll pathways (Scoffoni *et al*., 2014; Johnson *et al*., 2018). The reduced water potential in the xylem vessels leads to increased embolism and disruption of leaf hydraulic conductance, and ultimately the loss of rehydration capacity (Venturas *et al*., 2017; John *et al*., 2018). More recently, studies have shown that with severe dehydration there is loss of cellular function and viability which ultimately results in cell death and leaf necrosis (Trueba *et al*., 2019; Mantova *et al*., 2022). The sequence of breakdown of these physiological processes with increasing dehydration is typically correlated across species; i.e., species tolerant to mild dehydration are usually also tolerant to moderate and severe dehydration. This suggests coordination due to correlated selection or underlying mechanistic linkages (Bartlett *et al*., 2016; Trueba *et al*., 2019; Jin *et al*., 2023; Ziegler *et al*., 2024). The leaf water status that results in the breakdown of key physiological process, e.g. the relative water content (RWC) at loss of turgor, represents important species-specific thresholds that are related to drought tolerance (Anderegg *et al*., 2016; Torres-Ruiz *et al*., 2024). Thus, quantification of species thresholds has allowed assessment of vulnerabilities of species, forests and ecosystems to drought (Maherali *et al*., 2004; Bartlett *et al*., 2016).

The substantial progress in identifying the important physiological processes that are disrupted with increasing dehydration is in contrast to the general lack of studies that have examined the dynamics of water loss during drought. Plants continue to lose water even after complete stomatal closure (Schreiber & Riederer, 1996; Schuster *et al*., 2017). This residual water loss, leaf minimum conductance (*g*_min_), occurs via the cuticle, or through leaky stomata (Duursma *et al*., 2019). The magnitude of water loss via *g*_min_ can be substantial, and can vary considerably across species (Schreiber & Riederer, 1996; Schuster *et al*., 2017; Duursma *et al*., 2019; Ochoa *et al*., 2024). Such estimates of *g*_min_ are important for quantifying water loss, water use efficiency, and gas exchange across scales from leaves to ecosystems (De Kauwe *et al*., 2020; Lanning *et al*., 2020; Ochoa *et al*., 2024). There is growing evidence to suggest that *g*_min_ is important in species responses to drought, and may determine the time to reach critical levels of dehydration (Gleason *et al*., 2014; Blackman *et al*., 2016, 2019, 2023; Martin-StPaul *et al*., 2017; De Kauwe *et al*., 2020; Machado *et al*., 2021; Wang *et al*., 2024; Ziegler *et al*., 2024). However, our understanding of how important *g*_min_ is in determining the time species take to reach critical levels of dehydration, how it is related to other hydraulic traits that contribute to drought tolerance, and the determinants of variation in *g*_min_ across species is still incomplete (Choat *et al*., 2018; Brodribb *et al*., 2020; Blackman *et al*., 2023; Ochoa *et al*., 2024).

Physiological thresholds like the water potential that results in a 50 % disruption of hydraulic conductance are widely used as indices of drought tolerance, and species that can maintain function at lower levels of dehydration are assumed to be more tolerant (Bartlett *et al*., 2016; Trueba *et al*., 2019; Jin *et al*., 2023; Ziegler *et al*., 2024). However, in solely using such physiological thresholds as a measure of drought tolerance, an implicit assumption being made is that differences in water loss between species are negligible. Avoiding dehydration by minimizing water loss can represent an alternate strategy to cope with drought (Martin-StPaul *et al*., 2017), and the rates of residual water loss, and differences between species can be substantial (Schreiber & Riederer, 1996; Schuster *et al*., 2017; Duursma *et al*., 2019). Thus, overall tolerance to drought may be better represented by measures that incorporate both species thresholds and the rates of water loss (Blackman et al., 2016). The time taken by species when exposed to drought to reach critical thresholds that result in impairment of function represents such an integrated measure. Results from studies that have examined how *g*_min_ is related to thresholds for hydraulic dysfunction suggest these may not be related (Petek-Petrik *et al*., 2023; Waite *et al*., 2024), or may even be positively related to each other (Martin-StPaul *et al*., 2017; Ziegler *et al*., 2024). Thus, species with high drought tolerance may also have high *g*_min_, indicative of a tradeoff between tolerance (thresholds) and avoidance (water loss via *g*_min_) (Martin-StPaul *et al*., 2017). As a consequence, the rank order of species drought tolerance based on thresholds, may not reflect the ranks based on the time taken by species to reach these physiological thresholds.

To understand how leaf minimum conductance (*g*_min_) is related to critical physiological thresholds, we quantified relative water content (RWC) based thresholds for early, moderate and severe dehydration. We selected the RWC that resulted in turgor loss, which is mechanistically linked to stomatal closure (Bartlett *et al*., 2016; Martin-StPaul *et al*., 2017), as the threshold for early dehydration. Turgor loss point thresholds vary widely (Maréchaux *et al*., 2015), and are related to species responses to drought (McGregor *et al*., 2021), and species distributions across gradients of water availability (Baltzer *et al*., 2008). We used the RWC that caused significant leaf shrinkage, an indicator of the loss of leaf structural integrity and disruption in leaf hydraulic conductance (Scoffoni *et al*., 2014, 2017) as a threshold related to moderate dehydration. Leaf shrinkage is associated with the breakdown of the extra-xylem hydraulic pathways, and the loss of cell-cell connectivity (Sancho-Knapik *et al*., 2011; Scoffoni *et al*., 2014; Buckley, 2015). Finally, we used the leaf RWC that results in the breakdown in photosystem-II (PSII) function as a threshold for severe dehydration. Recent studies have suggested that the tolerance of leaf photochemistry to dehydration is a promising trait to assess plant drought tolerance (Fortunel *et al*., 2023). The breakdown of PSII function is directly linked to cell viability, cellular death and leaf necrosis (Mantova *et al*., 2022). Unlike turgor loss and shrinkage, which represent thresholds from which leaves can potentially recover, disruption of PSII function represents irreversible damage that ultimately leads to cell death (Mantova *et al*., 2021).

We used leaf dehydration assays to quantify leaf minimum conductance (*g*_min_) in 18 coexisting evergreen and deciduous trees from a seasonally dry tropical forest in peninsular India. To determine tolerance to different intensities of water stress from early to severe dehydration, we quantified the RWC based thresholds for loss of turgor, breakdown of structural integrity, and disruption of cellular function in leaves of these species. In the course of the leaf dehydration assays we quantified the time taken by species to reach these critical physiological breakpoints. This allowed us to test how time to reach thresholds was related to *g*_min_ and the thresholds themselves. Additionally, we quantified stomatal traits, key leaf functional traits, and other hydraulic traits to test how these are related to leaf minimum conductance (*g*_min_). Finally, we used a principal component analysis to determine the major axes of variation and hence strategies that emerge from the traits examined.

## Materials and Methods

### Site description, species selection and sample collection

The Northern Western Ghats region in peninsular India is highly seasonal, and most of the mean annual precipitation of around 2266 mm occurs during the monsoon months between June and October (New *et al*., 2002). The long dry season with average monthly rainfall less than 100 mm extends from November to May. The landscape is topographically diverse with valleys carved out by the Bhima river and its tributaries. The top of the valleys consist of flattened ridges with low soil depth and high light availability, and are characterized by vegetation of low stature and a higher percentage of deciduous species (open forests). In contrast, the bottom of the valleys that have greater soil depth and low light availability in the understorey, are dominated by tall statured evergreen species (closed forests). The transition vegetation in the slopes of the valleys are intermediate between the closed and open forests.

Evergreen and deciduous leaf habits that represent distinct strategies to cope with the seasonally varying water availability are both common in our study site (Sunny *et al*., 2025). Unlike deciduous species that shed their leaves in the dry season, evergreen species typically minimize water loss and maximize drought tolerance to maintain their canopy throughout the year (Álvarez-Yépiz *et al*., 2017). We selected 18 angiosperm tree species (11 evergreen and 7 deciduous) based on their dominance in a seasonally dry tropical forest near Nigdale, Maharashtra, India. The cumulative basal area of these 18 species is greater than 80 % of the total basal area in these forests (Jazeera *et al*., 2016). Mature individuals of each species were selected, and upper canopy sun-exposed branches of around 1 m in length were collected using a combination of tree-climbers and telescopic tree pruners.

### Leaf dehydration assay and quantification of leaf minimum conductance

For 6 replicate individuals of each species, 2^nd^ or 3^rd^ order branches containing fully expanded and mature leaves were collected between March and May of 2017. These branches were placed in a darkened plastic bag, and the bags were sealed with moist paper towels to keep the air in the bag water-saturated for transportation to the laboratory. On the same day in the laboratory, the leaves were cut underwater, placed in beakers with petioles immersed in water, and put in a sealed plastic bag in the dark for overnight rehydration (> 12 h). The next morning, saturated fresh weight (SFW) was quantified for one mature leaf from each individual. The leaf was then kept upright and allowed to dry in the laboratory on a drying rack at low irradiance. A table fan set at low speed was used to minimize boundary layer conductance throughout the assay. Temperature and humidity was monitored using a custom-built Arduino-based data logger and sensors. The average relative humidity in the room was 38.1 ± 7.0 %, and the average temperature was 27.1 ± 2.1 °C. After the initial saturated fresh weight measurement, the leaf was weighed after the 1^st^, 2^nd^, 3^rd^, 6^th^, 9^th^ and 13^th^ hour, and subsequently, after every 12 hours of drying. The assay continued till the dark-adapted fluorescence measurements (described in the next section) reached zero, indicating a complete loss of cellular function. The leaves were subsequently oven-dried for 72 hours at 70 °C, and the dry weight (DW) quantified. The relative water content (RWC) corresponding to every time point (*t*) for a given leaf was calculated as:

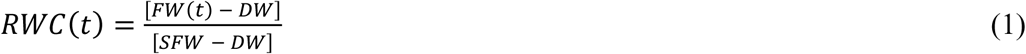

where FW(*t*) corresponds to the leaf weight at time *t*. An exponential decay model was fit to RWC as a function of time, and this was used to determine the time at which a leaf lost 50 % of its RWC (Time_RWC50_).

Leaf minimum conductance (*g*_min_) was determined using protocols described by Sack et al. (2003) and Sack and Scoffoni (2010). Briefly, this was quantified as the rate of water loss from the leaf after stomatal closure, divided by the vapour pressure deficit (VPD) and normalized by the double-sided leaf area. The first hour of drying was excluded, to account for complete stomatal closure, and the slope of the subsequent linear portion of the leaf drying curve was used to quantify *g*_min_. Leaf area was measured using a desktop scanner and ImageJ (version 1.47v; National Institutes of Health, Maryland, US).

### Leaf shrinkage

The same leaf used for the above assay was used to quantify leaf thickness and shrinkage using a digital micrometer (±0.002 mm, Mitutoyo) during dehydration (Scoffoni *et al*., 2014). The initial thickness was measured for the water saturated leaf before the initiation of drying, and then subsequently after the 1^st^, 2^nd^, 3^rd^, 6^th^, 9^th^ and 13^th^ hour of drying, and subsequently, after every 12 hours. The final leaf thickness was measured for the oven-dried leaves. Leaf thickness was normalized by the initial thickness measured for the water-saturated leaf to calculate relative thickness. An exponential decay model was fit to the pooled data from all individuals of a species for change in relative thickness as a function of leaf RWC. The leaf RWC that corresponded to a 50 % decrease in relative leaf thickness was determined (RWC_shrink50_), and used as a measure of leaf tolerance to shrinkage during dehydration. The time taken to lose 50 % leaf thickness during dehydration (Time_shrink50_) was determined for each species by fitting a three-parameter exponential decay model to relative leaf thickness as a function of time.

### Chlorophyll a fluorescence

The maximum quantum yield of photosystem II (PSII) was quantified as the ratio of dark-adapted variable fluorescence to maximum fluorescence (*F_v_/F_m_*), where *F_v_* = *F_m_* − *F_0_*, and *F_m_* and *F_0_* are the maximum and basal fluorescence, respectively. These Chlorophyll *a* fluorescence measurements were made on the same leaves used for the previous assays. Leaves were dark-adapted for 20 minutes, and fluorescence measurements made on the adaxial side, towards the center of the leaf avoiding the midrib, with a PAM 2500 fluorometer (Walz, Effeltrich, Germany). The maximum quantum yield of PSII is an index of leaf photochemical performance, and 50 % loss of PSII function is indicative of irreversible breakdown of the photosynthetic machinery and cellular function. A five-parameter logistic model was fit to the *F_v_/F_m_* response to RWC, and the RWC that resulted in the initial 5 % decrease (RWC_flbrk_), and the 50 % decrease (RWC_fl50_) in *F_v_/F_m_* was determined. The time taken to lose 5 % and 50 % of PSII function during dehydration (Time_flbrk_ and Time_fl50_) were determined for each species by fitting a five-parameter logistic model for *F_v_/F_m_* as a function of time.

### Pressure-volume curves to determine tolerance to turgor loss

A second bench drying assay for generating leaf pressure-volume curves was carried out during November and December in 2017 and 2018. Sun-exposed, upper canopy leaves were collected from five individuals of every species, rehydrated overnight as described above, and subjected to drying. The leaves were weighed, and water potential (PMS pressure chamber, Model 1515D) quantified at intervals of 0.2 to 0.3 MPa till a water potential of −3 MPa was reached. The leaves were subsequently oven-dried at 70 °C for 72 hours to determine the dry weight, and the RWC estimated (as described above). The pressure-volume curves were used to estimate water potential at the turgor loss point (*Ψ*_TLP_), RWC at the turgor loss point (RWC_TLP_), modulus of elasticity (*e*), and capacitance (CFT), using the protocol described by Sack et al. (2011). We used RWC_TLP_ for each species generated from the pressure-volume curves to estimate the time to loss of turgidity (Time_TLP_) from the species-level exponential decay model corresponding to loss of leaf RWC as a function of time, obtained from the assay described above.

### Stomatal traits

Six mature and sun-exposed leaves were collected from five of the same individuals of each study species. Nail polish varnish imprints were obtained from the central portion of the abaxial side taking care to avoid the midrib and major veins. Stomatal density, guard cell length, and pore length were quantified from images of the imprints using ImageJ (Schneider *et al*., 2012). Stomatal density (StomDen) was estimated as the number of stomata in a 20 × magnified field of view, and stomatal pore length (Pore size) and guard cell length (GCL) measured from 40 × magnified images. The maximum stomatal conductance (*g*_wmax_) for water was calculated using the following equation (Franks & Farquhar, 2001; Franks *et al*., 2009):

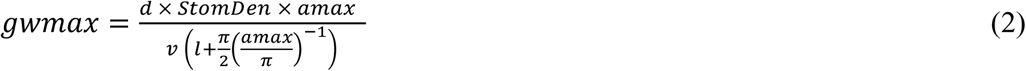

where *d* is the diffusivity of water vapour in air (2.49 × 10^-5^ m^2^·s^-1^), *amax* is the maximum stomatal pore area (m^−2^) calculated as an ellipse of area π × (0.5 × pore length) × (0.25 × pore length), v is molar volume of air (0.0224 m^3^·mol^−1^), and *l* is the depth of the stomatal pore which was approximated as 0.25 × Guard cell length. We could not obtain proper peels for two of the 18 study species. For *Callicarpa tomentosa*, stomatal peels were patchy due to the presence of leaf hair, and this allowed estimation of dimensions for individual stomata, but not stomatal density. For *Mallotus phillipensis*, neither stomatal dimensions nor density could be quantified, but data from a published study (Gangadhara, 2016) were used for our analyses.

### Leaf functional traits

A minimum of five mature and sun-exposed leaves were collected from each of the same six individuals of every species for quantifying leaf area, leaf dry matter content (LDMC) and leaf mass per area (LMA) (Pérez-Harguindeguy *et al*., 2013). Leaves were rehydrated overnight, as described above. Saturated fresh weight and leaf area were quantified for the rehydrated leaves, and dry weight after oven drying at 70 °C for 72 hours. LDMC was calculated as leaf dry weight per saturated fresh weight (g·g^-1^), and LMA as leaf dry weight per leaf area (g·m^-2^).

### In-situ leaf water status and phenology

To measure the leaf water status of the species in the driest time of the year, sun-exposed upper canopy leaves were collected at the end of the summer between 1230 and 1500 hours in May 2018. Leaf RWC was quantified for five or more mature leaves from 6 replicate individuals of each species. The leaves were individually stored in previously weighed, darkened, and sealed plastic bags for transport to the laboratory and was weighed within three hours of collection. The leaves were then rehydrated overnight by immersing the petioles in a beaker of water. The saturated fresh weight (SFW) of the rehydrated leaves was measured the following day, and the leaves were oven-dried at 70 °C for 72 hours before measurement of their dry weight (DW). RWC was calculated as in equation (1) to estimate the in-situ minimum relative water content (RWC_min_).

Midday leaf water potential at the driest time of the year (*Ψ*_min_) was measured at the end of the dry season in April and May of 2019 for 12 of the 18 study species. *Ψ*_min_ for the remaining 5 species were quantified in April and May of 2024. We were unable to determine *Ψ*_min_ for *Ficus racemosa* due to the non-availability of healthy leaves during the dry season. Five mature leaves were collected from three replicate individuals of each species between 1230 and 1500 hours from the sun-exposed upper canopy. Water potential was measured immediately after collection using a portable pressure chamber (Model 1515D, PMS Instruments Co., Albany, OR, USA).

In addition to the discrete evergreen and deciduous leaf habit categories, we examined the relationship of *g*_min_ and physiological thresholds to quantitative measures of leaf phenology. Woody species in these seasonal forests vary in how much of their canopy they lose in the dry season, and also in the duration that they remain leafless (Chakrabarty *et al*., 2021). The total canopy of individuals was scored by direct visual observation in a semi-quantitative manner on a scale from 0 to 100 % in steps of 10 %. A score of zero represented the complete absence of leaves, while a score of 100 represented a full canopy. Observations were conducted monthly for four years between 2014 and 2017. The average canopy maintained by individuals over the year was estimated, and 100 − average canopy, a measure of the average canopy loss (ACL), was used as a quantitative measure of deciduousness.

### Data Analysis

We used the ‘drc’ package (Ritz & Streibig, 2005) in R (version 4.1.1) for fitting logistic functions to the dehydration responses, and quantification of the corresponding thresholds (RWC_TLP_, RWC_shrink50_, RWC_flbrk_, RWC_fl50_), and the time to reach the thresholds (Time_RWC50_, Time_shrink50_, Time_TLP_, Time_flbrk_, Time_fl50_). All measured and derived traits were tested for normality using the Shapiro-Wilk test and transformed when necessary. We examined variation in traits using nested analysis of variance, with species nested within leaf habit. Bivariate trait relationships were assessed using Spearman’s rank correlation analysis. Multivariate trait dimensions were assessed using a principal component analysis (PCA) with the ‘factoextra’ package (Kassambara & Mundt, 2020) in R (version 4.1.1). We could not include stomatal traits, or RWC_min_ in the PCA analysis as we did not have data for all of the study species. All analyses were conducted using R (version 4.1.1).

**Table 1:** The traits examined, abbreviations and units.

|  | Traits | Abbreviation | Unit |
| --- | --- | --- | --- |
| a) Rate of water loss | Minimum cuticular conductance | $g_{min}$ | $mmol \cdot m^{-2} \cdot s^{-1}$ |
| b) Critical thresholds | RWC at Turgor Loss | $RWC_{TLP}$ | % |
| | RWC at 50 % leaf thickness | $RWC_{shrink50}$ | % |
| | RWC at 5 % decrease in $F_v/F_m$ | $RWC_{flbrk}$ | % |
| | RWC at 50 % decrease in $F_v/F_m$ | $RWC_{fl50}$ | % |
| c) Time to critical thresholds | Time to $RWC_{50}$ | $Time_{RWC50}$ | Hours |
| | Time to $RWC_{TLP}$ | $Time_{TLP}$ | Hours |
| | Time $RWC_{shrink50}$ | $Time_{shrink50}$ | Hours |
| | Time to $RWC_{flbrk}$ | $Time_{flbrk}$ | Hours |
| | Time to $RWC_{fl50}$ | $Time_{fl50}$ | Hours |
| d) Stomatal traits | Pore size | Pore size | $\mu m$ |
| | Guard cell length | GCL | $\mu m$ |
| | Stomatal density | StomDen | $mm^{-2}$ |
| | Max. stomatal conductance – water | $g_{wmax}$ | $mol \cdot m^{-2} \cdot s^{-1}$ |
| e) Other hydraulic traits | Water Potential at Turgor Loss | $\Psi_{TLP}$ | MPa |
| | Leaf capacitance at full turgor | CFT | $mol \cdot m^{-2} \cdot MPa^{-1}$ |
| | Modulus of elasticity at full turgor | $E$ | MPa |
| g) Morphological structural traits | Leaf area | LA | $cm^2$ |
| | Leaf dry matter content | LDMC | $g \cdot g^{-1}$ |
| | Leaf mass per area | LMA | $g \cdot m^{-2}$ |
| h) *In situ* traits | Average canopy loss | ACL | % |
| | Minimum relative water content | $RWC_{min}$ | % |
| | Minimum water potential | $\Psi_{min}$ | MPa |

## Results

There was large variation in the rate of water loss from leaves of the 18 study species, and the time to reach an RWC of 50 % varied more than 10-fold, ranging from around 2 to 21 hours (Fig. 1, Table S3). Deciduous species had higher rates of residual water loss, and reached 50 % RWC faster than evergreen species (Fig. 1, Table S3). Leaf minimum conductance (*g*_min_) varied more than six-fold ranging from 1.21 to 7.41 mmol·m^-2^·s^-1^, and was higher for deciduous than evergreen species (Fig. 1 inset, Table S3). The residual loss of water was not constant, and rates of water loss decreased with decreasing RWC. Methods to quantify *g*_min_ typically make measurements during the early hours of drying (Sack & Scoffoni, 2010; Slot *et al*., 2021), but for some species, like *Memecylon umbellatun*, there was still considerable water loss even on the third day of drying. To understand how *g*_min_ was related to water loss over longer periods of time, we examined how our estimates of *g*_min_ was related to time to reach RWC of 50 % (Time_RWC50_). There was a significant negative relationship between *g*_min_ and Time_RWC50_, indicating that species with higher residual water loss early during drying reached an RWC of 50 % sooner (Fig. S6).

**Figure 1:**
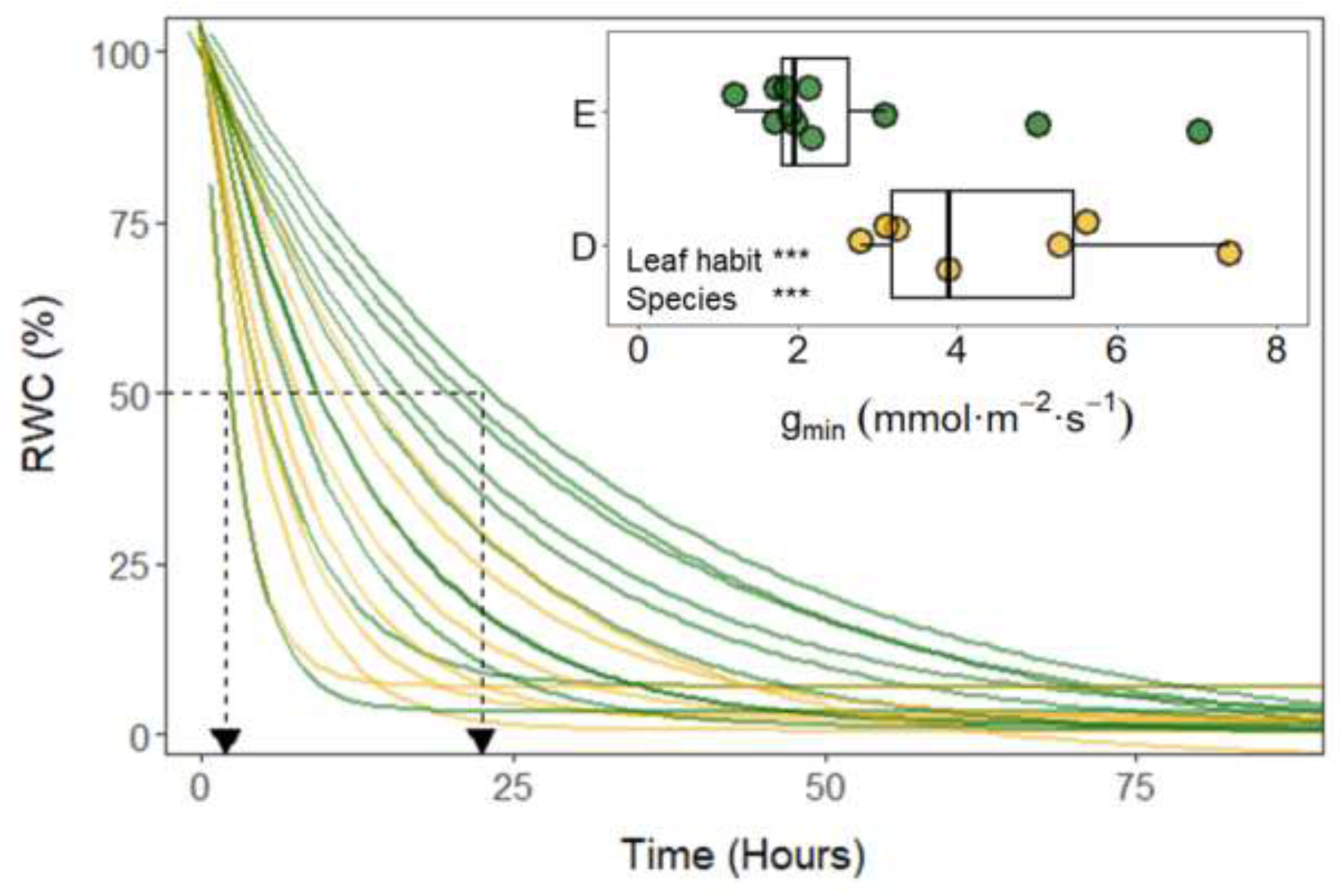
Loss of water quantified as change in RWC during leaf dehydration for the 18 study species (evergreen − green, deciduous − yellow). Dashed lines indicate the range of time taken to reach 50 % RWC for species with the highest and lowest rates of water loss. The inset shows box and whisker plots for leaf minimum conductance (*g*_min_) for the evergreen (E) and deciduous (D) species. ANOVA results for differences between leaf habit (evergreen and deciduous), and across species nested within leaf habit are depicted by *** for *P* < 0.01.

The RWC-based thresholds for early, moderate and severe dehydration differed across species (Fig. 2). The leaf RWC that resulted in loss of turgor (RWC_TLP_) ranged from 95 to 78.6 % across species (Fig. 2a, d, Table S3). The water potential at turgor loss (*Ψ*_TLP_) ranged from −0.59 to - 2.51 MPa MPa (Table S3), and was not related to RWC_TLP_ (r = 0.30, *p* = 0.21). The loss of leaf structural integrity (RWC_shrink50_) occurred at a lower RWC than turgor loss (for all but one of the species examined), and ranged from 90. 4 to 70.8 % (Fig. 2b, d, Table S3). Breakdown of PSII function, indicative of loss of cellular function and viability, occurred at much lower leaf RWC for all species (Fig. 2c, d, Table S3). The RWC that resulted the initial breakdown of PSII function (RWC_flbrk_) ranged from 65.5 to 26.5 %, and for 50 % loss of PSII function (RWC_fl50_) from 33.3 to 7.6 %. RWC_flbrk_ was closely related to RWC_fl50_ (Fig. S7) and for all further analysis we use RWC_flbrk_ as the threshold for PSII breakdown. Importantly, we found no evidence of any relationships between species thresholds for loss of turgor, breakdown of structural integrity, and loss of cellular function and viability (Fig. S8). Additionally, there were no significant relationships between all three thresholds and *g*_min_ (Fig. S8).

**Figure 2:**
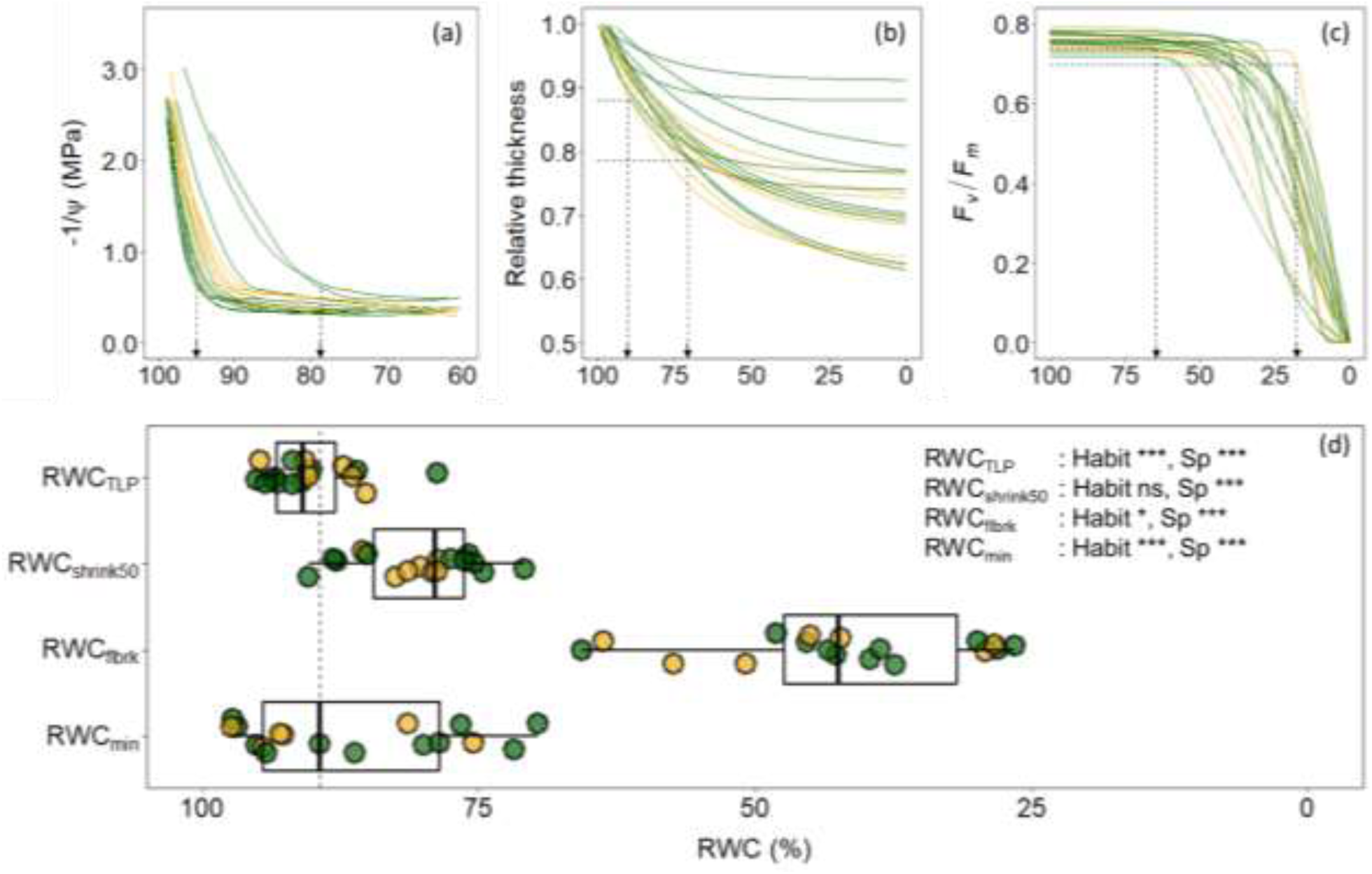
Pressure-volume curves, leaf thickness, and Photosystem II (PSII) function during leaf dehydration for the study species (evergreen − green, deciduous − yellow). These assays were used to determine the leaf RWC thresholds for: (a) turgor loss (RWC_TLP_); (b) loss of structural integrity (RWC_shrink50_); and, (c) breakdown of PSII function (RWC_flbrk_). Dashed lines indicates species with minimum and maximum threshold values. (d) Box-plots for these physiological thresholds, and minimum leaf RWC (RWC_min_). ANOVA results for differences between leaf habit (evergreen and deciduous), and across species nested within leaf habit are depicted by *** for *P* < 0.01, * for *P* < 0.1 and ‘ns’ for not significant.

The RWC threshold for turgor loss (RWC_TLP_) was lower for the deciduous than for the evergreen species, while in contrast, the threshold for breakdown in PSII function (RWC_flbrk_) was lower for the evergreen than for the deciduous species (Fig 2d). There was no difference between evergreen and deciduous species in the RWC thresholds for loss of structural integrity (RWC_shrink50_) (Fig 2d). The minimum leaf RWC (RWC_min_) for these species in their natural habitats at the driest time of the year differed across species, and was lower for the evergreen compared to the deciduous species (Fig. 2d, Table S3). RWC_min_ ranged from around to 97 to 69 % (Fig. 2d, Table S3), and for 10 of the 18 study species RWC_min_ was lower than RWC_TLP_. RWC_min_ was also lower than RWC_shrink50_ for 3 species, but higher than RWC_flbrk_ for all species.

The sequence of events with respect to time (Fig. 3) was similar to the sequence with respect to RWC (Fig. 2d). Turgor loss occurred rapidly, typically within 1 to 2 hours since the initiation of dehydration, and this did not differ between evergreen and deciduous species (Fig. 3). There was large variation across species in the time to 50 % shrinkage (Time_shrink50_) which ranged from 40 minutes to 15 hours, and in the time to loss PSII function (Time_flbrk_) which ranged from 2 hours to more than 36 hours. Leaves of deciduous species reached critical thresholds for loss of structural integrity and loss of PSII function earlier than the evergreen species (Fig. 3, Table S3).

**Figure 3:**
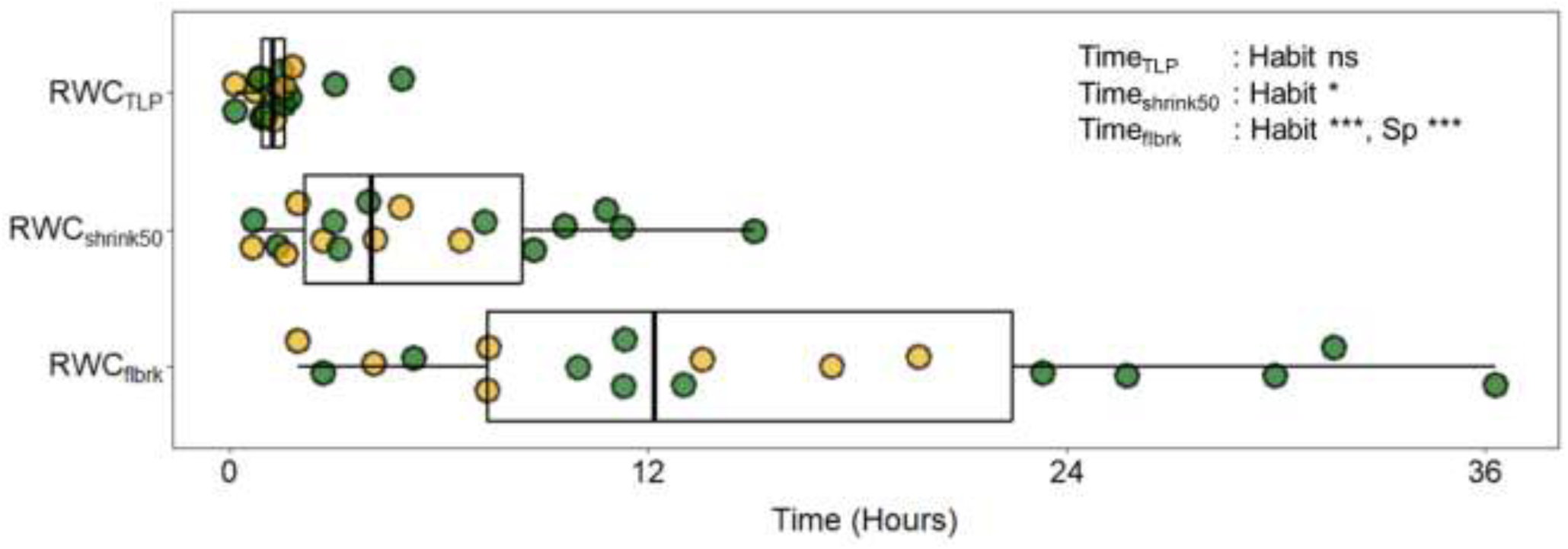
Time taken to reach the critical thresholds (RWC_TLP_, RWC_shrink50_, and RWC_flbrk_) for the 18 study species (yellow – deciduous, green – evergreen). ANOVA results for differences across leaf habit (evergreen and deciduous), and across species nested within leaf habit are depicted by *** for *P* < 0.01, * for *P* < 0.1 and ‘ns’ for not significant.

The time required for species to reach all three RWC thresholds was negatively related to *g*_min_ (Fig. 4a, c, e).The results for relationships between the time to reach these critical thresholds and species thresholds themselves were mixed: while significant for loss of structural integrity (Fig. 4d), this was not significant for turgor loss and breakdown of PSII function (Fig. 4b, f).

**Figure 4:**
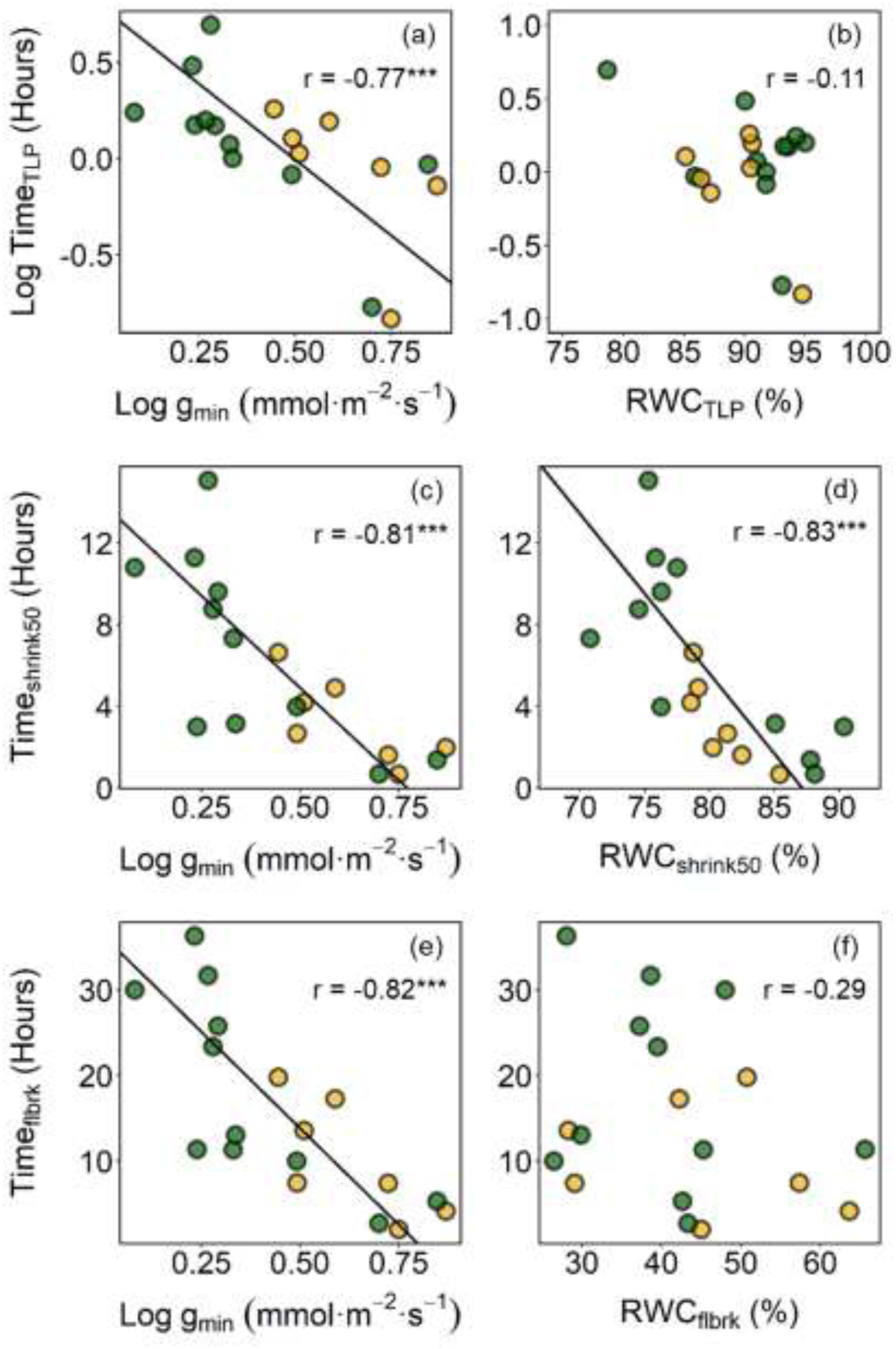
Relationship of the time taken to reach the thresholds with *g*_min_ and the thresholds themselves for the 18 study species (yellow – deciduous, green – evergreen). (a, b) time to turgor loss (log Time_TLP_); (c, d) Time to loss of structural integrity (Time_shrink50_); (e, f) Time to breakdown of PSII function (Time_flbrk_). Results for Spearman’s correlation coefficient (r) are shown, and solid lines represent type II regression fits for significant relationships (*P* < 0.01).

Leaf minimum conductance was positively related to stomatal density, and maximum stomatal conductance, but not related to pore size (Fig. S9). None of the other hydraulic, or leaf functional traits examined were related to *g*_min_ (Fig. S10). Finally, for the *in situ* parameters examined for these species in the field, *g*_min_ was positively related to the average canopy loss of species, but unrelated to the minimum RWC or minimum water potentials experienced by species in their natural environments during the driest time of the year (Fig. S10).

Results from the PCA showed that the first three PC dimensions explained more than 70 % of the total variation, with 31.5%, 26.4 %, and 12.7 % explained by the first, second and third PC axes, respectively (Fig. 5, Table S11). PC1 was primarily associated with rates of water loss and time to reach thresholds. PC1 was negatively correlated with time to loss of PSII function (Time_flbrk_) and structural integrity (Time_shrink50_), and positively correlated with *g*_min_ (Fig. 5, Table S12). This axis also had significant, but lower contributions from average canopy loss (ACL), the RWC threshold for structural integrity (RWC_shrink50_), time to turgor loss (Time_TLP_), and leaf mass per area (LMA). PC2 was primarily associated with species thresholds for early dehydration, and was positively related leaf capacitance (CFT) and time to turgor loss (Time_TLP_), and negatively related to leaf modulus of elasticity (*E*), the RWC threshold for turgor loss (RWC_TLP_), and leaf dry matter content (LDMC). Finally, PC3 was positively correlated with tolerance to severe dehydration (RWC_flbrk_) and leaf size (LA), and negatively correlated with water potential at turgor Loss (*Ψ*_TLP_).

**Figure 5:**
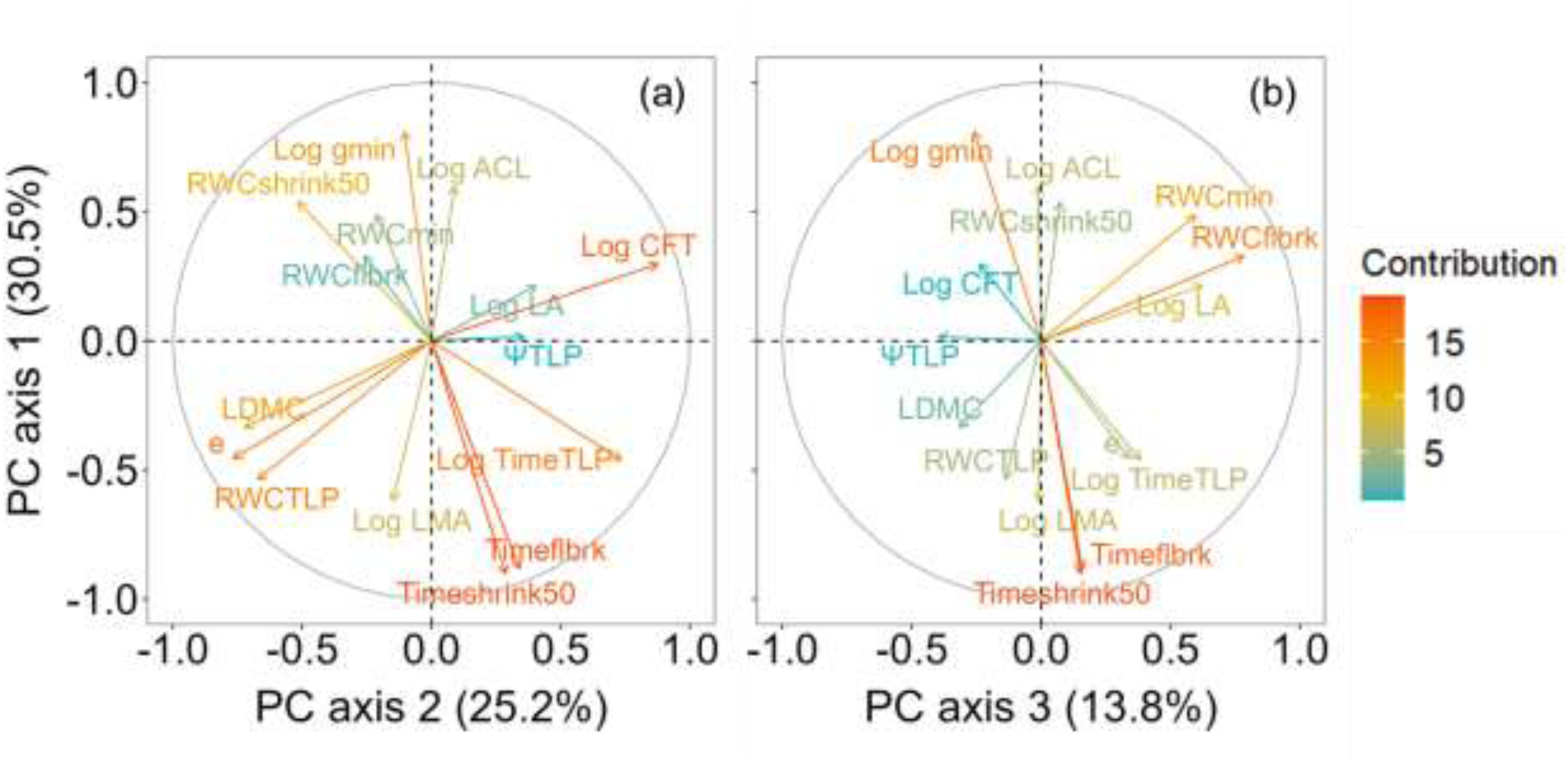
Principal component (PC) analysis of the leaf traits examined for the study species. Variable correlation plots for: (a) PC axes 1 and 2; and, (b) PC axes 2 and 3. The colour for the trait vectors represent the percent contributions to dimensions examined. Trait abbreviations and details are provided in Table 1.

## Discussion

We report significant variation in leaf minimum conductance (*g*_min_) across 18 coexisting trees from a seasonally dry tropical forest. Leaf minimum conductance varied 6-fold, and was greater for deciduous than evergreen species. Critical physiological thresholds indicative of tolerance to early, moderate and severe dehydration also differed across species, but were not related to each other, or to *g*_min_. Importantly, we found that *g*_min_ was the primary determinant of the time species took to reach these physiological thresholds. Results from a PCA analysis revealed three major independent axes of variation in hydraulic traits: The first represented variation in traits related to *g*_min_ and the time to reach critical thresholds; the second, tolerance to mild and moderate dehydration; and, the third, tolerance to severe dehydration. Surprisingly, *g*_min_ was negatively related to maximum stomatal conductance, suggesting that species can concurrently maximize conductance and minimize water loss via leaf minimum conductance.

Tropical species were poorly represented in previous global analysis of *g*_min_ (Schuster *et al*., 2017; Duursma *et al*., 2019), but there has been a recent increase in studies reporting *g*_min_ for tropical trees (Machado *et al*., 2021; Slot *et al*., 2021; Levionnois *et al*., 2021; Loram-Lourenço *et al*., 2022; Manzi *et al*., 2022; Ziegler *et al*., 2024; Wittemann *et al*., 2024; Middleby *et al*., 2024; Boisseaux *et al*., 2025). We collated this, and previous data for *g*_min_ of tropical species (> 200 species from > 8 sites) to contextualize our results. The range of *g*_min_ observed across our 18 species was similar to what was reported from a seasonal tropical forest in Brazil (Machado *et al*., 2021), but lower than the overall range across the tropics (Fig. S13). The higher *g*_min_ in deciduous than in evergreen species in our site is consistent with differences reported in most other tropical sites. While residual water loss represents a small fraction of the maximum potential leaf conductance, integrated over the cumulative leaf surface area for an individual, and over time of exposure to drought, the observed differences in *g*_min_ are likely to be significant and important. Minimizing water loss with reduced *g*_min_ may be particularly relevant for seasonally dry tropical forests like our study site, where species have to endure a harsh and prolonged dry season, and in future climates with higher temperatures and increased vapour pressure deficits (Hammond *et al*., 2022; Song *et al*., 2025).

As expected, loss of turgor and breakdown of structural integrity occurred at mild to moderate levels of dehydration, while breakdown of PSII function occurred with more severe dehydration at lower leaf RWC. The RWC threshold for turgor loss (RWC_TLP_) was lower for deciduous than evergreen species, indicating that deciduous species could sustain turgor at lower RWC levels and were more tolerant to mild dehydration. This would allow the more acquisitive deciduous species to continue carbon assimilation during early dehydration. In contrast, deciduous species had higher RWC thresholds for breakdown in PSII function (RWC_flbrk_). Thus, evergreen species that maintained most of their canopy through the dry season were more tolerant to severe dehydration levels that result in loss of cellular function and viability. While these physiological thresholds differed across species, these were not related to each other. Species that were more tolerant to mild dehydration were not necessarily the most tolerant to loss of structural integrity, or cell function and viability. This suggests lack of underlying mechanistic linkages, physiological constraints, or correlated selection for these traits in these species, unlike what has been reported for other hydraulic traits (Bartlett *et al*., 2016; Trueba *et al*., 2019).

The time taken by species to reach critical levels of dehydration integrates species-specific thresholds and rates of residual water loss. While turgor loss happened rapidly, there was large variation across species in the time to loss of structural integrity, and loss of PSII function. Interestingly, the time to these critical thresholds was primarily determined by *g*_min_. Thus, species with higher *g*_min_ reached critical thresholds for turgor loss, structural integrity loss, and PSII function breakdown earlier. Given that minimizing water loss and increasing physiological tolerance to dehydration can represent distinct, independent strategies to deal with drought, our results highlight the importance of moving beyond the current safety margin metrics to assess vulnerability of species to drought (Volaire, 2018; Trugman *et al*., 2021; McDowell *et al*., 2022). Such current safety margin metrics are exclusively based on physiological thresholds, whereas using integrated measures such as ‘time to physiological threshold’ may better capture species vulnerabilities. Not surprisingly, the traditional threshold-based safety margins (RWC_TLP_ − RWC_flbrk_) were not correlated with the time-based safety margin (Time_TLP_ − Time_flbrk_) for our study species (Fig. S10, S14). Our study, along with a growing body of literature (Blackman *et al*., 2016, 2023; Petek-Petrik *et al*., 2023; Ziegler *et al*., 2024; Burlett *et al*., 2025), calls attention to the need for such integrated measures of plant drought sensitivity.

Congruent with the lack of relationships between thresholds, and between thresholds and *g*_min_, the principle components analyses of trait combinations in these species showed multiple orthogonal axes of variation in hydraulic traits in our study species. The primary axis (PC1) captured variation associated with traits related to rates of residual water loss. This axis represented a spectrum with conservative evergreen species with low *g*_min_ and high LMA that were able to maintain structural integrity and cellular function for longer durations during dehydration at one end, and acquisitive deciduous species with high *g*_min_ and low LMA that lost structural integrity and cellular function earlier during dehydration at the other extreme. The second axis (PC2) represented a spectrum of variation in capacitance, LDMC, and tolerance to mild and moderate dehydration. Finally, the third axis (PC3) primarily represented variation in tolerance to severe dehydration, leaf area, and *in situ* minimum RWC experienced by species in their natural environment. Overall, our results suggest that these traits can evolve independently to allow a wider diversity of adaptive strategies to cope with the environmental conditions that vary in the severity of drought (Martin-StPaul *et al*., 2017; Blackman *et al*., 2019). In coexisting species, this would allow the fine tuning of strategies to balance opportunities for carbon gain with costs of potential breakdown in physiological function under varying intensities and durations of drought. The lack of coordination across measures of tolerance has important implications for ecologists and modelers, for extrapolating drought responses for a wide range of conditions based on a smaller number of traits (Bartlett *et al*., 2016; Kim *et al*., 2024).

A positive relationship of *g*_min_ with stomatal size and density may be expected if residual water loss occurs via incompletely closed or leaky stomata (Muchow & Sinclair, 1989; Ochoa *et al*., 2024). In addition to the mechanistic insights that such a relationship provides, this has important implications because increased stomatal size and density can drive higher stomatal conductance and enable greater carbon assimilation (Franks *et al*., 2009). A recent study found positive relationships between *g*_min_ and both stomatal density and carbon assimilation (Machado *et al*., 2021). This suggests a tradeoff where increased assimilation facilitated by higher conductance comes at the cost of greater water loss from leaky stomata. However, only one other study (Muchow & Sinclair, 1989) has reported a positive relationships between *g*_min_ and stomatal density, and the majority of others that have examined this report no relationships (summarized in Ochoa *et al*., 2024). Surprisingly, we observed a significant negative relationships between *g*_min_ and stomatal density, and *g*_min_ and maximum potential stomatal conductance. Thus, species with low residual water loss had higher stomatal densities and maximum stomatal conductance. This is congruent with results from causal analyses of stomatal traits and *g*_min_ by Ochoa et al. (2025), and confirm their inference that residual water loss may evolve independently of stomatal traits allowing species to simultaneously increase maximum stomatal conductance and minimize residual water loss.

The lack of a tradeoff with maximum stomatal conductance, and no clear relationships with other hydraulic and functional traits left open the question of why species maintained high residual water loss when they could minimize *g*_min_ at no apparent cost. One possible explanation could lie in the potential benefit that *g*_min_ can provide for maintaining safe operating leaf temperatures during drought. Droughts are frequently accompanied by high temperatures (IPCC 2021), and in the seasonal forests in our study site the hottest time of the year coincides with the late dry season (Sastry & Barua, 2018, Javad et al. 2025). Limited water availability and closed stomata may limit the ability of leaves to effectively cool themselves under these conditions (Evans *et al*., 2025). Recent studies have shown that the relative contribution of cuticular versus stomatal conductance increases dramatically at higher temperatures (Slot *et al*., 2021; Zailaa *et al*., 2024; Fernandes *et al*., 2025; Garen & Michaletz, 2025), and this may be important in cooling leaves exposed to extreme temperatures during drought. The temperature dependence of *g*_min_ indicates that the importance of residual water loss in driving impairment of leaf and plant function will be more dominant in future climates with hotter and more intense droughts.

## Conclusion

These results go beyond recent studies which suggest that residual water loss from leaves is important in determining the time to ultimate hydraulic failure and mortality (Blackman *et al*., 2016; Ziegler *et al*., 2024; Burlett *et al*., 2025). We show that *g*_min_ is central in determining the time taken to reach physiological thresholds that impair leaf function across a wide range of leaf water status from mild to severe dehydration. Differences in *g*_min_ and time to critical physiological thresholds in coexisting species can result in differential carbon assimilation, growth and survival in the face of varying intensities and duration of drought. Such *g*_min_ driven differences in plant performance are likely to be even more important in future climates with hotter climate change-type droughts. The independence of *g*_min_ with tolerance to mild and severe tolerance further emphasizes the importance of quantifying leaf minimum conductance in addition to other traits that contribute to drought tolerance to comprehensively assess vulnerability of species to drought.

## Acknowledgements

The authors wish to thank Kalu and Ganpath for assistance in field work; Dhrubojyoti Patra and Omkar Khache for help with sample collection and processing; Dr. Narendra Kadoo for providing the pressure chamber; Akhil Javad for help with the leaf economic budget model analysis; Indian Institute of Science Education and Research (IISER), Pune for intra-mural funds and support for Ron Sunny (Integrated PhD fellowship).

## Author contributions

RS and DB designed the study. RS collected the data with help from MV for the bench drying assays, and BP for quantification of stomatal traits. SC collected the phenology data. RS analyzed the data with help from DB. RS and DB wrote the initial drafts of the manuscript with input from MV, BP and SC.

## Data availability

The data that supports the findings of this study are available in the supplementary material of this article.

## References

Álvarez-Yépiz JC, Búrquez A, Martínez-Yrízar A, Teece M, Yépez EA, Dovciak M. 2017. Resource partitioning by evergreen and deciduous species in a tropical dry forest. Oecologia 183: 607–618.

Anderegg WRL, Klein T, Bartlett M, Sack L, Pellegrini AFA, Choat B, Jansen S. 2016. Meta-analysis reveals that hydraulic traits explain cross-species patterns of drought-induced tree mortality across the globe. Proceedings of the National Academy of Sciences 113: 5024–5029.

Baltzer JL, Davies SJ, Bunyavejchewin S, Noor NSM. 2008. The role of desiccation tolerance in determining tree species distributions along the Malay–Thai Peninsula. Functional Ecology 22: 221–231.

Bartlett MK, Klein T, Jansen S, Choat B, Sack L. 2016. The correlations and sequence of plant stomatal, hydraulic, and wilting responses to drought. Proceedings of the National Academy of Sciences 113: 13098–13103.

Blackman CJ, Billon L-M, Cartailler J, Torres-Ruiz JM, Cochard H. 2023. Key hydraulic traits control the dynamics of plant dehydration in four contrasting tree species during drought (T Holtta, Ed.). Tree Physiology 43: 1772–1783.

Blackman CJ, Brodribb TJ, Jordan GJ. 2010. Leaf hydraulic vulnerability is related to conduit dimensions and drought resistance across a diverse range of woody angiosperms. New Phytologist 188: 1113–1123.

Blackman CJ, Li X, Choat B, Rymer PD, De Kauwe MG, Duursma RA, Tissue DT, Medlyn BE. 2019. Desiccation time during drought is highly predictable across species of *Eucalyptus* from contrasting climates. New Phytologist 224: 632–643.

Blackman CJ, Pfautsch S, Choat B, Delzon S, Gleason SM, Duursma RA. 2016. Toward an index of desiccation time to tree mortality under drought: Desiccation time to tree mortality. Plant, Cell & Environment 39: 2342–2345.

Boisseaux M, Nemetschek D, Baraloto C, Burban B, Casado-Garcia A, Cazal J, Clément J, Derroire G, Fortunel C, Goret J, et al. 2025. Shifting trait coordination along a soil-moisture-nutrient gradient in tropical forests. Functional Ecology 39: 21–37.

Brodribb TJ, Powers J, Cochard H, Choat B. 2020. Hanging by a thread? Forests and drought. Science 368: 261–266.

Buckley TN. 2015. The contributions of apoplastic, symplastic and gas phase pathways for water transport outside the bundle sheath in leaves: Leaf water pathways. *Plant*, Cell & Environment 38: 7–22.

Burlett R, Trueba S, Bouteiller XP, Forget G, Torres-Ruiz JM, Martin-StPaul NK, Parise C, Cochard H, Delzon S. 2025. Minimum leaf conductance during drought: unravelling its variability and impact on plant survival. New Phytologist 246: 1001–1014.

Chakrabarty S, Sharma S, Ganguly S, Jezeera A, Mohanbabu N, Barua D. 2021. Quantitative estimates of deciduousness in woody species from a seasonally dry tropical forest are related to leaf functional traits and the timing of leaf flush. BioRxiv 2021.03.03: 433407 [preprint].

Choat B, Brodribb TJ, Brodersen CR, Duursma RA, López R, Medlyn BE. 2018. Triggers of tree mortality under drought. Nature 558: 531–539.

De Kauwe MG, Medlyn BE, Ukkola AM, Mu M, Sabot MEB, Pitman AJ, Meir P, Cernusak LA, Rifai SW, Choat B et al. 2020. Identifying areas at risk of drought-induced tree mortality across South-Eastern Australia. Global Change Biology 26:5716–5733.

Duursma RA, Blackman CJ, Lopéz R, Martin-StPaul NK, Cochard H, Medlyn BE. 2019. On the minimum leaf conductance: its role in models of plant water use, and ecological and environmental controls. New Phytologist 221: 693–705.

Evans MEK, Hu J, Michaletz ST. 2025. Scaling plant responses to heat: From molecules to the biosphere. Science 388(6752): 1167–1173.

Fernandes VDAB, Farnese FS, Arantes BR, Fontineles Da Silva ML, Silva FG, Torres-Ruiz JM, Slot M, Cochard H, Menezes-Silva PE. 2025. Leaf minimum conductance dynamics during and after heat stress: Implications for plant survival under hotter droughts. Plant Physiology 197: kiaf026.

Fortunel C, Stahl C, Coste S, Ziegler C, Derroire G, Levionnois S, Maréchaux I, Bonal D, Hérault B, Wagner FH, et al. 2023. Thresholds for persistent leaf photochemical damage predict plant drought resilience in a tropical rainforest. New Phytologist 239: 576–591.

Franks PJ, Drake PL, Beerling DJ. 2009. Plasticity in maximum stomatal conductance constrained by negative correlation between stomatal size and density: an analysis using *Eucalyptus globulus*. Plant, Cell & Environment 32: 1737–1748.

Franks PJ, Farquhar GD. 2001. The Effect of Exogenous Abscisic Acid on Stomatal Development, Stomatal Mechanics, and Leaf Gas Exchange in *Tradescantia virginiana*. Plant Physiology 125: 935–942.

Gangadhara M. 2016. Epidermal structure, structure and ontogeny of stomata in some dicotyledons. PhD. Thesis. Sardar Patel University, Gujarat, India.

Garen JC, Michaletz ST. 2025. Temperature governs the relative contributions of cuticle and stomata to leaf minimum conductance. New Phytologist 245: 1911–1923.

Gleason SM, Blackman CJ, Cook AM, Laws CA, Westoby M. 2014. Whole-plant capacitance, embolism resistance and slow transpiration rates all contribute to longer desiccation times in woody angiosperms from arid and wet habitats. Tree Physiology 34: 275–284.

Hammond WM, Adams HD. 2019. Dying on time: traits influencing the dynamics of tree mortality risk from drought. Tree Physiology 39: 906–909.

Hammond WM, Williams AP, Abatzoglou JT, Adams HD, Klein T, López R, Sáenz-Romero C, Hartmann H, Breshears DD, Allen CD. 2022. Global field observations of tree die-off reveal hotter-drought fingerprint for Earth’s forests. Nature Communications 13: 1761.

Intergovernmental Panel on Climate Change (IPCC). 2021. In: VP Masson-Delmotte et al., eds. Climate Change 2021: The Physical Science Basis. Working Group I contribution to the Sixth Assessment Report of the Intergovernmental Panel on Climate Change. Geneva, Switzerland: IPCC.

Javad A, Premugh V, Tiwari R, Bandaru P, Sunny R, Hegde B, Clerici S, Galbraith D, Gloor M, Barua D. 2025. Leaf temperatures in an indian tropical forest exceed physiological limits but durations of exposures are currently not sufficient to cause lasting damage. Global Change Biology, 31: p.e70069.

Jezeera, AM. 2016. Variation in plant functional traits across contrasting habitats in a seasonally dry tropical forest in the Northern Western Ghats. Master’s Thesis Dissertation. Indian Institute of Science Education and Reseacrch, Pune, India.

Jin Y, Hao G, Hammond WM, Yu K, Liu X, Ye Q, Zhou Z, Wang C. 2023. Aridity-dependent sequence of water potentials for stomatal closure and hydraulic dysfunctions in woody plants. Global Change Biology 29: 2030–2040.

John GP, Henry C, Sack L. 2018. Leaf rehydration capacity: Associations with other indices of drought tolerance and environment. Plant, Cell & Environment 41: 2638–2653.

Johnson KM, Jordan GJ, Brodribb TJ. 2018. Wheat leaves embolized by water stress do not recover function upon rewatering. *Plant*, Cell & Environment 41: 2704–2714.

Kassambara A, Fabian M. 2020. Factoextra: Extract and Visualize the Results of Multivariate Data Analyses. https://cran.r-project.org/web/packages/factoextra/index.html.

Kim D, Guadagno CR, Ewers BE, Mackay DS. 2024. Combining PSII photochemistry and hydraulics improves predictions of photosynthesis and water use from mild to lethal drought. Plant, Cell & Environment 47: 1255–1268.

Klein T, Yakir D, Buchmann N, Grünzweig JM. 2014. Towards an advanced assessment of the hydrological vulnerability of forests to climate change-induced drought. New Phytologist 201: 712–716.

Lanning M, Wang L, Novick KA. 2020. The importance of cuticular permeance in assessing plant water–use strategies. Tree Physiology 40: 425–432.

Levionnois S, Ziegler C, Heuret P, Jansen S, Stahl C, Calvet E, Goret J-Y, Bonal D, Coste S. 2021. Is vulnerability segmentation at the leaf-stem transition a drought resistance mechanism? A theoretical test with a trait-based model for Neotropical canopy tree species. Annals of Forest Science 78: 87.

Loram-Lourenço L, Farnese FS, Alves RDFB, et al. 2022. Variations in bark structural properties affect both water loss and carbon economics in neotropical savanna trees in the Cerrado region of Brazil. Journal of Ecology 110: 1826–1843

Machado R, Loram-Lourenço L, Farnese FS, Alves RDFB, de Sousa LF, Silva FG, Filho SCV, Torres-Ruiz JM, Cochard H, Menezes-Silva PE. 2021. Where do leaf water leaks come from? Trade-offs underlying the variability in minimum conductance across tropical savanna species with contrasting growth strategies. New Phytologist 229: 1415–1430.

Maherali H, Pockman WT, Jackson RB. 2004. Adaptive variation in the vulnerability of woody plants to xylem cavitation. Ecology 85: 2184–2199.

Mantova M, Herbette S, Cochard H, Torres-Ruiz JM. 2022. Hydraulic failure and tree mortality: from correlation to causation. Trends in Plant Science 27: 335–345.

Mantova M, Menezes-Silva PE, Badel E, Cochard H, Torres-Ruiz JM. 2021. The interplay of hydraulic failure and cell vitality explains tree capacity to recover from drought. Physiologia Plantarum 172: 247–257.

Manzi OJL, Bellifa M, Ziegler C, Mihle L, Levionnois S, Burban B, Leroy C, Coste S, Stahl C. 2022. Drought stress recovery of hydraulic and photochemical processes in Neotropical tree saplings (I Ensminger, Ed.). Tree Physiology 42: 114–129.

Maréchaux I, Bartlett MK, Sack L, Baraloto C, Engel J, Joetzjer E, Chave J. 2015. Drought tolerance as predicted by leaf water potential at turgor loss point varies strongly across species within an Amazonian forest (K Kitajima, Ed.). Functional Ecology 29: 1268–1277.

Martinez-Vilalta J, Anderegg WRL, Sapes G, Sala A. 2019. Greater focus on water pools may improve our ability to understand and anticipate drought-induced mortality in plants. New Phytologist 223: 22–32.

Martin-StPaul N, Delzon S, Cochard H. 2017. Plant resistance to drought depends on timely stomatal closure (H Maherali, Ed.). Ecology Letters 20: 1437–1447.

McDowell NG, Sapes G, Pivovaroff A, Adams HD, Allen CD, Anderegg WRL, Arend M, Breshears DD, Brodribb T, Choat B, et al. 2022. Mechanisms of woody-plant mortality under rising drought, CO2 and vapour pressure deficit. Nature Reviews Earth & Environment 3: 294– 308.

McGregor IR, Helcoski R, Kunert N, Tepley AJ, Gonzalez-Akre EB, Herrmann V, Zailaa J, Stovall AEL, Bourg NA, McShea WJ, et al. 2021. Tree height and leaf drought tolerance traits shape growth responses across droughts in a temperate broadleaf forest. New Phytologist 231: 601–616.

Middleby KB, Cheesman AW, Cernusak LA. 2024. Impacts of elevated temperature and vapour pressure deficit on leaf gas exchange and plant growth across six tropical rainforest tree species. New Phytologist 243: 648–661.

Muchow RC, Sinclair TR. 1989. Epidermal conductance, stomatal density and stomatal size among genotypes of *Sorghum bicolor* (L.) Moench. Plant, Cell & Environment 12: 425–431.

New M, Lister D, Hulme M, Makin I. 2002. A high-resolution data set of surface climate over global land areas. Climate Research 21: 1–25.

Ochoa ME, Henry C, John GP, Medeiros CD, Pan R, Scoffoni C, Buckley TN, Sack L. 2024. Pinpointing the causal influences of stomatal anatomy and behavior on minimum, operational, and maximum leaf surface conductance. Plant Physiology 196: 51–66.

Pérez-Harguindeguy N, Díaz S, Garnier E, Lavorel S, Poorter H, Jaureguiberry P, Bret-Harte MS, Cornwell WK, Craine JM, Gurvich DE, et al. 2013. New handbook for standardised measurement of plant functional traits worldwide. Australian Journal of Botany 61: 167.

Petek-Petrik A, Petrík P, Lamarque LJ, Cochard H, Burlett R, Delzon S. 2023. Drought survival in conifer species is related to the time required to cross the stomatal safety margin (T Lawson, Ed.). Journal of Experimental Botany 74: 6847–6859.

Powers JS, Vargas-G G, Brodribb TJ, Schwartz NB, Perez-Aviles D, Smith-Martin CM, Becknell JM, Aureli F, Blanco R, Calderón-Morales E, et al. 2020. A catastrophic tropical drought kills hydraulically vulnerable tree species. Global Change Biology 26: 3122–3133..

Ritz C, Streibig JC. 2005. Bioassay Analysis using *R*. Journal of Statistical Software 12: 1–22.

Sack L, Cowan PD, Jaikumar N, Holbrook NM. 2003. The ‘hydrology’ of leaves: co-ordination of structure and function in temperate woody species. Plant Cell and Environment 26:1343–1356.

Sack L, Scoffoni C. 2010. Minimum epidermal conductance (*g*_min_ a.k.a. cuticular conductance). Prometheus Wiki.

Sack L, Pasquet-Kok J, Bartlett M. 2010. Leaf pressure-volume curve parameters. Prometheus Wiki.

Sancho-Knapik D, Álvarez-Arenas TG, Peguero-Pina JJ, Fernández V, Gil-Pelegrín E. 2011. Relationship between ultrasonic properties and structural changes in the mesophyll during leaf dehydration. Journal of Experimental Botany 62: 3637–3645.

Sastry A, Barua D. 2017. Leaf thermotolerance in tropical trees from a seasonally dry climate varies along the slow-fast resource acquisition spectrum. Scientific Reports 7:11246.

Schreiber L, Riederer M. 1996. Ecophysiology of cuticular transpiration: comparative investigation of cuticular water permeability of plant species from different habitats. Oecologia 107: 426–432.

Schneider CA, Rasband WS, Eliceiri KW. 2012. NIH Image to ImageJ: 25 years of image analysis. Nature Methods 9: 671–675

Schuster A-C, Burghardt M, Riederer M. 2017. The ecophysiology of leaf cuticular transpiration: are cuticular water permeabilities adapted to ecological conditions? Journal of Experimental Botany 68: 5271–5279.

Scoffoni C, Albuquerque C, Brodersen CR, Townes SV, John GP, Bartlett MK, Buckley TN, McElrone AJ, Sack L. 2017. Outside-Xylem Vulnerability, Not Xylem Embolism, Controls Leaf Hydraulic Decline during Dehydration. Plant Physiology 173: 1197–1210.

Scoffoni C, Vuong C, Diep S, Cochard H, Sack L. 2014. Leaf Shrinkage with Dehydration: Coordination with Hydraulic Vulnerability and Drought Tolerance. Plant Physiology 164: 1772– 1788.

Slot M, Nardwattanawong T, Hernández GG, Bueno A, Riederer M, Winter K. 2021. Large differences in leaf cuticle conductance and its temperature response among 24 tropical tree species from across a rainfall gradient. New Phytologist 232: 1618–1631.

Song F, Dong H, Wu L, Leung LR, Lu J, Dong L, Wu P, Zhou T. 2025. Hot season gets hotter due to rainfall delay over tropical land in a warming climate. Nature Communications 16: 2188.

Sunny R, Guha A, Jezeera A, Mohan N K, Mohanbabu N, Barua D. 2025. Responses to water limitation are independent of light for saplings of a seasonally dry tropical forest. Biotropica 57: e13404.

Torres-Ruiz JM, Cochard H, Delzon S, Boivin T, Burlett R, Cailleret M, Corso D, Delmas CEL, Caceres M, Diaz-Espejo A et al. 2024. Plant hydraulics at the heart of plant, crops and ecosystem functions in the face of climate change. New Phytologist 241: 984–999.

Trueba S, Pan R, Scoffoni C, John GP, Davis SD, Sack L. 2019. Thresholds for leaf damage due to dehydration: declines of hydraulic function, stomatal conductance and cellular integrity precede those for photochemistry. New Phytologist 223: 134–149.

Trugman AT, Anderegg LDL, Anderegg WRL, Das AJ, Stephenson NL. 2021. Why is Tree Drought Mortality so Hard to Predict? Trends in Ecology & Evolution 36: 520–532.

Venturas MD, Sperry JS, Hacke UG. 2017. Plant xylem hydraulics: What we understand, current research, and future challenges. Journal of Integrative Plant Biology 59: 356–389.

Volaire F. 2018. A unified framework of plant adaptive strategies to drought: Crossing scales and disciplines. Global Change Biology 24: 2929–2938.

Wang S, Hoch G, Grun G, Kahmen A. 2024. Water loss after stomatal closure: quantifying leaf minimum conductance and minimal water use in nine temperate European tree species during a severe drought. Tree Physiology 44: tpae027.

Waite PA, Kumar M, Link RM, Schuldt B. 2024. Coordinated hydraulic traits influence the two phases of time to hydraulic failure in five temperate tree species differing in stomatal stringency. Tree Physiology 44: tpae038.

Wittemann M, Mujawamariya M, Ntirugulirwa B, Uwizeye FK, Zibera E, Manzi OJL, Nsabimana D, Wallin G, Uddling J. 2024. Plasticity and implications of water-use traits in contrasting tropical tree species under climate change. Physiologia Plantarum 176: e14326.

Zailaa J, Scoffoni C, Brodersen CR. 2024. Stomatal closure as a driver of minimum leaf conductance declines at high temperature and vapor pressure deficit in *Quercus*. Plant Physiology 197: kiae551.

Ziegler C, Cochard H, Stahl C, Foltzer L, Gérard B, Goret J-Y, Heuret P, Levionnois S, Maillard P, Bonal D, et al. 2024. Residual water losses mediate the trade-off between growth and drought survival across saplings of 12 tropical rainforest tree species with contrasting hydraulic strategies (M Moshelion, Ed.). Journal of Experimental Botany: erae159.

