## supplemental info for "It’s about time! Leaf minimum conductance determines time to reach critical thresholds for leaf dehydration in a seasonal tropical forest"

### Supplementary figures and tables

**Table S1:** Species list and trait values

**Table S2:** Species list and traits values (continued)

**Table S3:** F statistic for ANOVA for effect of leaf habit (evergreen and deciduous) and species (nested in leaf habit) on the traits

**Figure S1:** Map showing study site

**Figure S2:** Species level curves showing time to leaf dehydration

**Figure S3:** Species level pressure volume curves

**Figure S4:** Dehydration response curves of the leaf relative thickness for each species

**Figure S5:** Dehydration response curves for leaf PSII function ($F_v/F_m$) for each species

**Figure S6:** Relation between $g_{min}$ and $Time_{RWC50}$

**Figure S7:** Relation between $RWC_{flbrk}$ and $RWC_{fl50}$

**Figure S8:** Relation between the resistance to loss of turgidity, structural integrity and cellular function and rate of leaf water loss

**Figure S9:** Relation between $g_{min}$ and stomatal traits

**Figure S10:** Spearman's correlation coefficient of the traits examined in the study

**Figure S11:** Scree plot of the principal component analysis

**Figure S12:** Contributions of variables in PCA components

**Figure S13:** Literature review summary of $g_{min}$

**Figure S14:** Schematic concept figure

**Table S1:** Species, taxonomic affiliation, species codes (ID), and trait values for the 18 study species: $g_{min}$ ($mmol \cdot m^{-2} \cdot s^{-1}$), $RWC_{TLP}$ (%), $\Psi_{TLP}$ (MPa), $RWC_{shrink50}$ (%), $RWC_{fl50}$ (%), $RWC_{flbrk}$ (%), $Time_{RWC50}$ (Hours), $Time_{TLP}$ (Hours), $Time_{shrink50}$ (Hours), $Time_{flbrk}$ (Hours) and $Time_{fl50}$ (Hours). Trait abbreviations are provided in Table 1.

| | **Species** | **Family** | **ID** | $g_{min}$ | **RWC TLP** | **$\Psi$ TLP** | **RWC shrink50** | **RWC fl50** | **RWC flbrk** | **Time RWC50** | **Time TLP** | **Time shrink50** | **Time flbrk** | **Time fl50** |
|---|---|---|---|---|---|---|---|---|---|---|---|---|---|---|
| 1. | *Actinodaphne angustifolia* | Lauraceae | AC | 2.14 | 91.0 | -2.20 | 70.8 | 22.0 | 45.3 | 8.8 | 1.2 | 7.3 | 11.3 | 15.7 |
| 2. | *Atalantia racemosa* | Rutaceae | AR | 1.96 | 93.6 | -1.81 | 76.3 | 12.6 | 37.3 | 16.7 | 1.5 | 9.6 | 25.8 | 40.6 |
| 3. | *Bridelia retusa* | Phyllanthaceae | BR | 7.41 | 87.2 | -1.73 | 80.3 | 33.3 | 63.6 | 3.6 | 0.7 | 2.0 | 4.1 | 5.4 |
| 4. | *Psydrax dicoccos* | Rubiaceae | CD | 1.71 | 90.0 | -2.29 | 75.8 | 18.1 | 28.1 | 20.2 | 3.0 | 11.3 | 36.3 | 46.1 |
| 5. | *Macaranga peltata* | Euphorbiaceae | CH | 7.03 | 85.9 | -1.72 | 87.8 | 28.9 | 42.7 | 4.5 | 0.9 | 1.4 | 5.3 | 7.8 |
| 6. | *Catunaregam Spinosa* | Rubiaceae | CS | 5.29 | 86.4 | -2.02 | 82.5 | 13.2 | 29.1 | 4.4 | 0.9 | 1.6 | 7.4 | 12.2 |
| 7. | *Callicarpa tomentosa* | Lamiaceae | CT | 1.90 | 78.7 | -1.70 | 74.5 | 14.2 | 39.6 | 15.3 | 4.9 | 8.7 | 23.3 | 31.9 |
| 8. | *Diospyros montana* | Ebenaceae | DM | 3.24 | 90.5 | -2.14 | 78.6 | 10.8 | 28.3 | 7.5 | 1.1 | 4.2 | 13.5 | 21.2 |
| 9. | *Flacourtia indica* | Salicaceae | FI | 3.11 | 85.1 | -2.51 | 81.4 | 18.2 | 57.4 | 5.7 | 1.3 | 2.7 | 7.4 | 13.6 |
| 10. | *Ficus racemosa* | Moraceae | FR | 5.62 | 94.8 | -1.29 | 85.4 | 25.0 | 45.0 | 2.4 | 0.1 | 0.7 | 2.0 | 4.7 |
| 11. | *Mangifera indica* | Anacardiaceae | MI | 1.73 | 93.3 | -1.92 | 90.4 | 32.7 | 65.6 | 12.6 | 1.5 | 3.0 | 11.3 | 19.2 |
| 12. | *Mallotus phillipensis* | Euphorbiaceae | MP | 5.02 | 93.1 | -2.19 | 88.1 | 14.4 | 43.4 | 2.2 | 0.2 | 0.7 | 2.7 | 4.5 |
| 13. | *Memecylon umbellatum* | Melastomataceae | MU | 1.85 | 95.0 | -1.67 | 75.3 | 12.5 | 38.6 | 21.6 | 1.6 | 15.1 | 31.7 | 50.4 |
| 14. | *Olea dioica* | Oleaceae | OD | 2.17 | 91.8 | -1.58 | 85.1 | 10.1 | 29.9 | 8.6 | 1.0 | 3.1 | 13.0 | 22.8 |
| 15. | *Syzygium cumini* | Myrtaceae | SC | 1.21 | 94.3 | -1.73 | 77.5 | 20.4 | 48.0 | 20.4 | 1.7 | 10.8 | 30.0 | 41.9 |
| 16. | *Terminalia bellirica* | Combretaceae | TB | 3.88 | 90.6 | -1.52 | 79.1 | 17.8 | 42.3 | 11.3 | 1.6 | 4.9 | 17.3 | 25.2 |
| 18. | *Terminalia chebula* | Combretaceae | TC | 2.79 | 90.4 | -1.61 | 78.7 | 19.9 | 50.8 | 12.8 | 1.8 | 6.6 | 19.7 | 25.6 |
| 19. | *Xantolis tomentosa* | Sapotaceae | XT | 3.10 | 91.8 | -0.59 | 76.3 | 7.6 | 26.5 | 6.8 | 0.8 | 4.0 | 10.0 | 16.0 |

**Table S2:** Species ID (details in table S3), leaf habit (LH: E – evergreen, D – deciduous), and trait values for the 18 study species. Stomatal pore size (Pore size, µm), guard cell length (GCL, µm), stomatal density (StomDen, $mm^{-2}$), maximum stomatal conductance ($g_{wmax}$, $mol \cdot m^{-2} \cdot s^{-1}$), leaf area (LA, $cm^2$), leaf dry matter content (LDMC, $g \cdot g^{-1}$), leaf mass per area (LMA, $g \cdot m^{-2}$), capacitance at full turgor (CFT, $mol \cdot m^{-2} \cdot MPa^{-1}$), cell wall rigidity (*e*, MPa), average canopy loss (ACL,%), minimum *in situ* RWC ($RWC_{min}$, %), minimum *in situ* water potential ($\Psi_{min}$, MPa).

| | **ID** | **LH** | **Pore size** | **GCL** | **StomDen** | $g_{wmax}$ | **LA** | **LDMC** | **LMA** | $CFT_{abs}$ | ***e*** | **ACL** | $RWC_{min}$ | $\Psi_{min}$ |
|---|---|---|---|---|---|---|---|---|---|---|---|---|---|---|
| 1. | AC | E | 13.0 | 20.3 | 268.7 | 1.62 | 44.8 | 0.462 | 119.1 | 0.358 | 20.6 | 3.24 | 95.1 | -1.60 |
| 2. | AR | E | 5.2 | 8.1 | 800.2 | 1.91 | 25.2 | 0.405 | 123.4 | 0.454 | 23.7 | 5.75 | 80.0 | -3.38 |
| 3. | BR | D | 10.3 | 14.0 | 195.2 | 0.98 | 42.1 | 0.358 | 117.6 | 0.897 | 12.9 | 12.53 | 92.6 | -1.57 |
| 4. | CD | E | 12.8 | 19.2 | 493.3 | 2.96 | 49.7 | 0.446 | 198.6 | 0.708 | 17.3 | 4.33 | 71.8 | -3.67 |
| 5. | CH | E | 9.4 | 15.4 | 254.2 | 1.08 | 133.0 | 0.391 | 118.2 | 0.837 | 10.4 | 4.52 | 96.8 | -0.67 |
| 6. | CS | D | 10.0 | 21.5 | 413.0 | 1.66 | 15.6 | 0.364 | 111.1 | 0.893 | 11.7 | 30.76 | 81.3 | -1.78 |
| 7. | CT | E | 11.3 | 19.7 | — | — | 126.3 | 0.282 | 73.3 | 2.277 | 5.2 | 5.94 | 78.5 | -1.68 |
| 8. | DM | D | 23.6 | 31.7 | 180.7 | 2.07 | 29.3 | 0.372 | 99.7 | 0.599 | 19.0 | 10.67 | 75.5 | -3.82 |
| 9. | FI | D | 14.3 | 22.0 | 1074.0 | 7.13 | 33.1 | 0.388 | 86.7 | 0.547 | 13.8 | 30.30 | 94.5 | -1.79 |
| 10. | FR | D | 14.3 | 19.5 | 313.7 | 2.19 | 43.2 | 0.363 | 115.2 | 0.408 | 19.4 | 24.80 | — | — |
| 11. | MI | E | 7.8 | 18.6 | 749.3 | 2.20 | 60.7 | 0.437 | 131.9 | 0.380 | 25.6 | 7.70 | 97.2 | -0.24 |
| 12. | MP | E | 7.7 | 13.0 | 496.0 | 1.70 | 31.8 | 0.472 | 88.0 | 0.232 | 24.9 | 6.21 | 89.4 | -1.67 |
| 13. | MU | E | 14.9 | 25.4 | 350.9 | 2.30 | 19.2 | 0.439 | 204.4 | 0.448 | 20.6 | 3.93 | 76.5 | -3.95 |
| 14. | OD | E | 20.2 | 27.0 | 276.2 | 2.71 | 34.7 | 0.443 | 123.4 | 0.456 | 16.0 | 5.79 | 69.6 | -4.64 |
| 15. | SC | E | 12.2 | 19.6 | 949.6 | 5.33 | 46.8 | 0.378 | 103.8 | 0.336 | 24.0 | 6.45 | 94.1 | -1.43 |
| 16. | TB | D | 21.3 | 29.0 | 268.5 | 2.75 | 79.3 | 0.331 | 142.5 | 1.024 | 14.6 | 11.18 | 97.3 | -1.23 |
| 17. | TC | D | 17.3 | 23.1 | 340.3 | 2.89 | 58.8 | 0.327 | 127.6 | 0.927 | 15.5 | 16.06 | 92.9 | -1.27 |
| 18. | XT | E | 16.3 | 19.6 | 183.0 | 1.51 | 25.1 | 0.442 | 98.0 | 2.141 | 4.4 | 7.24 | 86.2 | -3.31 |

**Table S3:** Differences in traits between leaf habit (evergreen and deciduous) and species. ANOVA results (F-statistics and significance) for leaf habit and species (nested within leaf habit). Some of the traits examined do not include results for species (nested within leaf habit) due to lack individual level data within species. Significance levels: $P < 0.01$ – ***; $P < 0.05$ – **, $P < 0.1$ – *; not significant – ns. Abbreviations for traits are described in Table 1.

| | Traits | Leaf habit | | Species \| Leaf habit | |
|---|---|---|---|---|---|
| a) Rate of water loss | $\log(g_{min})$ | 99.2 | *** | 14 | *** |
| b) Critical thresholds | $RWC_{TLP}$ | 17.3 | *** | 19.3 | *** |
| | $RWC_{shrink50}$ | 0.2 | ns | 7 | *** |
| | $RWC_{fl50}$ | 2.6 | ns | 9.9 | *** |
| | $RWC_{flbrk}$ | 3.6 | * | 4.4 | *** |
| c) Time thresholds | $Time_{RWC50}$ | 76.9 | *** | 15.3 | *** |
| | $\log(Time_{TLP})$ | 0.8 | ns | | |
| | $Time_{shrink50}$ | 3.6 | * | | |
| | $Time_{fl50}$ | 68.5 | *** | 18.5 | *** |
| | $Time_{flbrk}$ | 58.1 | *** | 17.1 | *** |
| d) Stomatal traits | Pore size | 327.6 | *** | 84.9 | *** |
| | Guard cell length | 233.5 | *** | 58.7 | *** |
| | Stomatal density | 46.3 | *** | 79 | *** |
| | $\log(g_{wmax})$ | 12 | *** | 44.2 | *** |
| e) Other hydraulic traits | $\Psi_{TLP}$ | 0.7 | ns | 18.2 | *** |
| | log(CFT) | 4.5 | ** | 8.6 | *** |
| | $E$ | 12.5 | *** | 11 | *** |
| g) Morphological functional traits | log(LA) | 16.9 | *** | 54.8 | *** |
| | LDMC | 87.7 | *** | 12.3 | *** |
| | log(LMA) | 7.1 | *** | 21.6 | *** |
| h) *In situ* traits | log(ACL) | 48.0 | *** | | |
| | $RWC_{min}$ | 13.7 | *** | 22.9 | *** |
| | $\Psi_{min}$ | 7.4 | ** | 9.6 | *** |

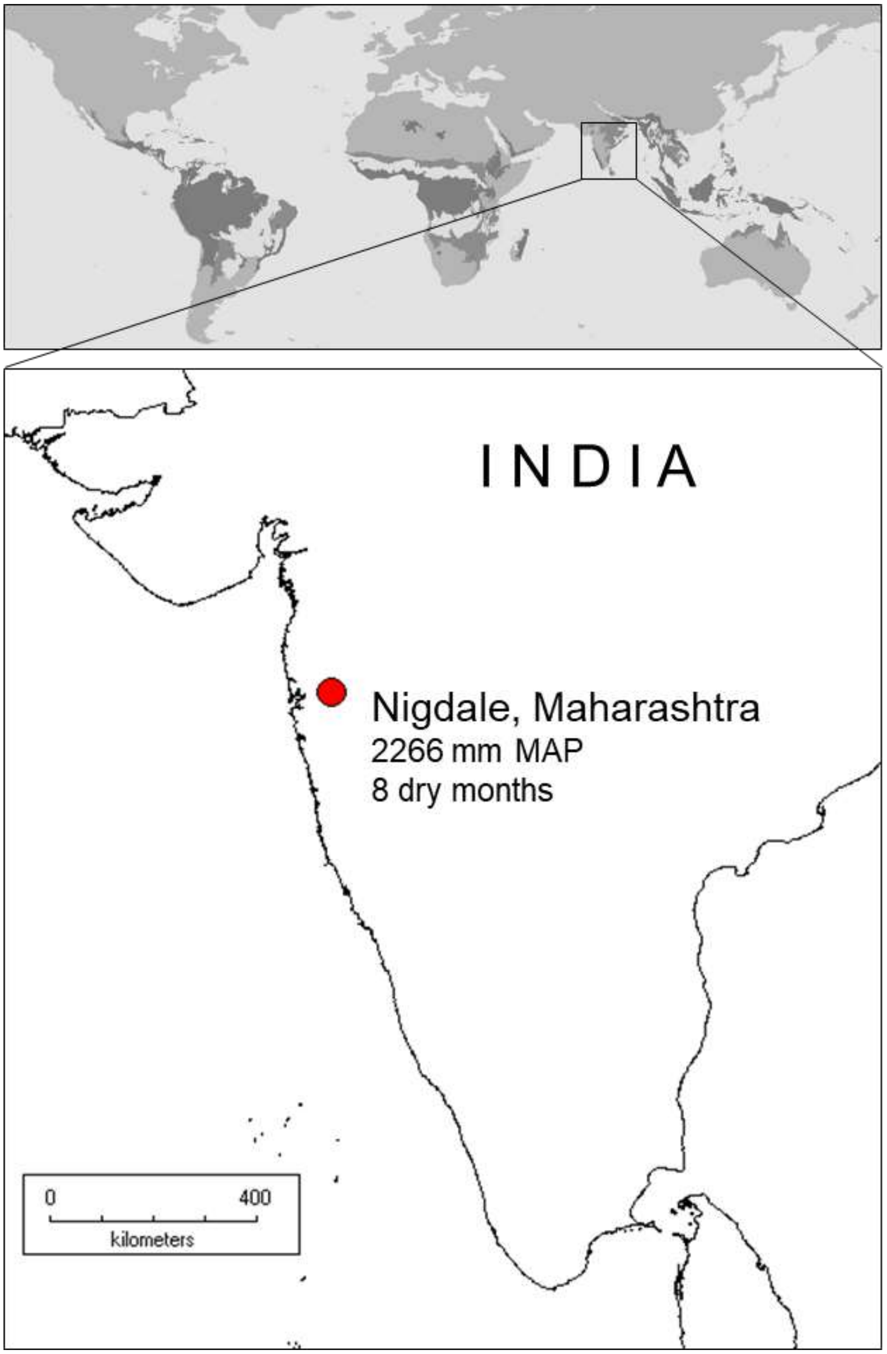


**Figure S1:** Map showing location of the seasonally dry tropical forest located in Nigdale, Maharashtra, in the northern part of the Northern Western Ghats in India. Eighteen dominant angiosperm adult tree species were sampled from this forest for the present study.

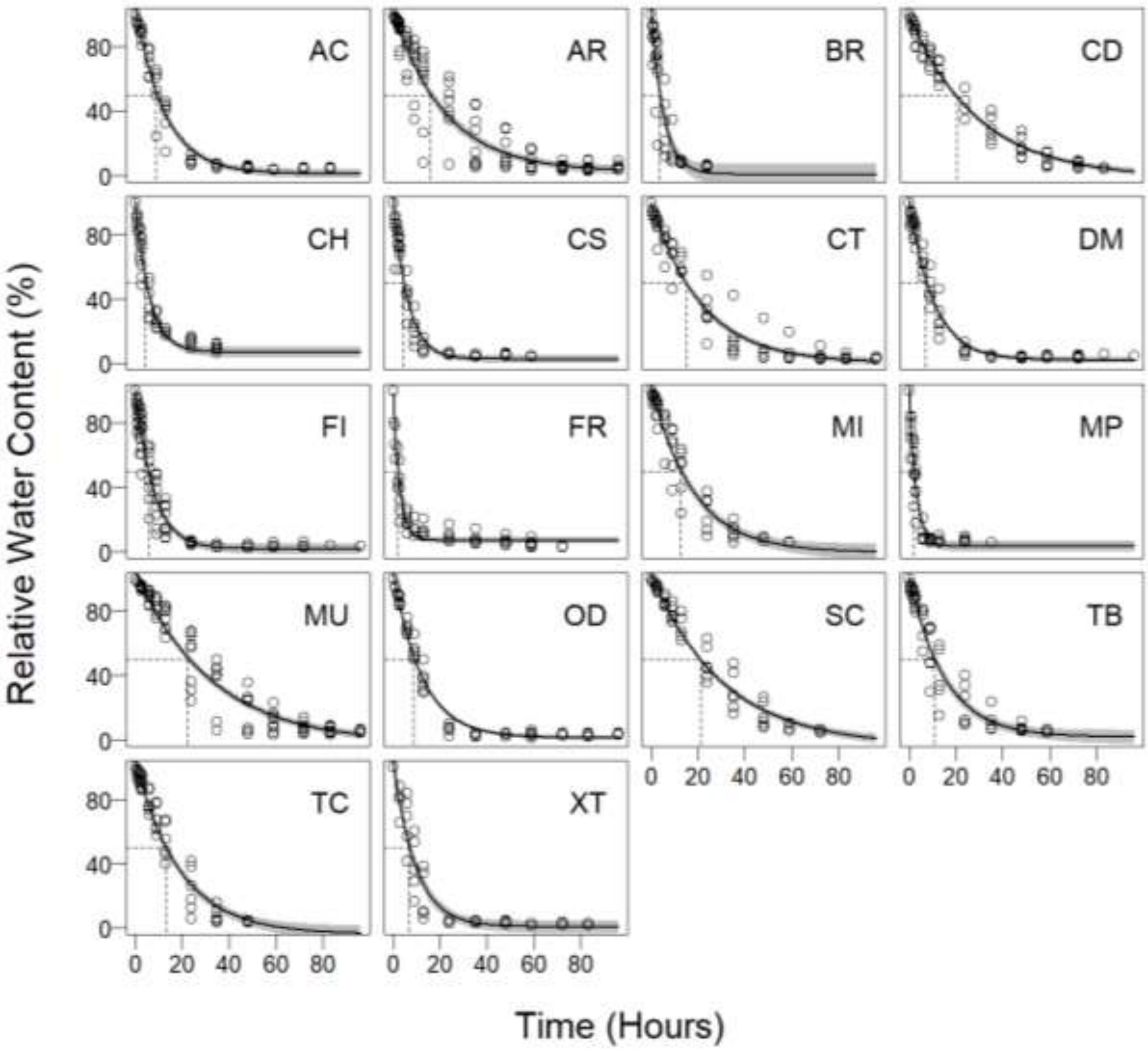


**Figure S2:** Time to leaf dehydration for the 18 species studied from its water saturated stage as represented by leaf relative water content lost over time in hours. Water status of the leaf in terms of the relative water content (RWC) ranges from 100% (well hydrated) to 0% (completely dry). Each graph is data pooled from at least 5 replicate individuals. A three parameter exponential decay function fit to the data is shown with grey confidence interval band. The line correspond to the time at which the species lost 50% of its leaf relative water content. The abbreviations for the species are as in table S1.

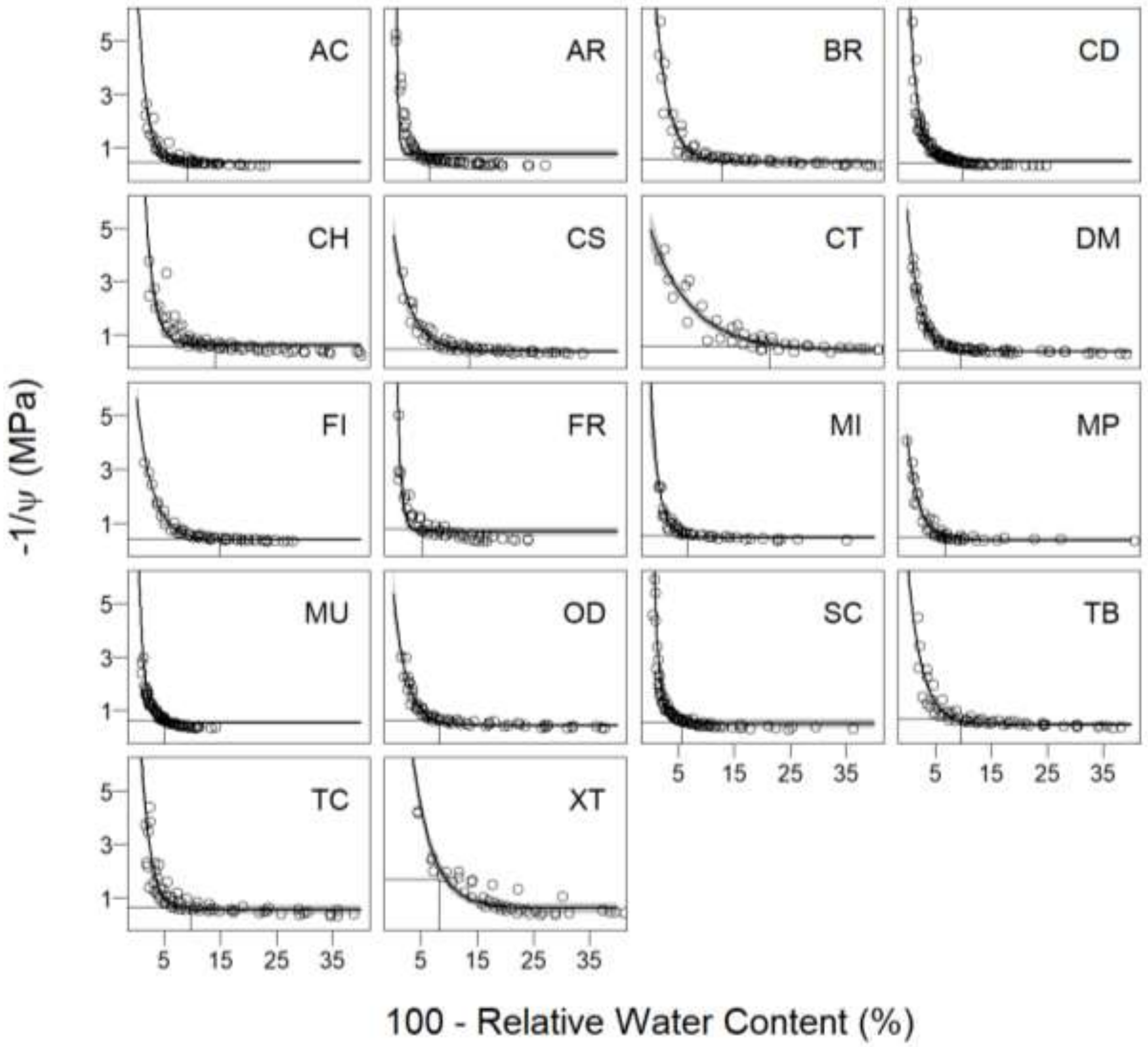


**Figure S3:** Pressure–volume curve data for the 18 species studied. Each graph is data pooled from at least 5 replicate individuals. The $RWC_{TLP}$ and the $\Psi_{TLP}$ for each of the species is shown. The abbreviations for the species are as in table S1.

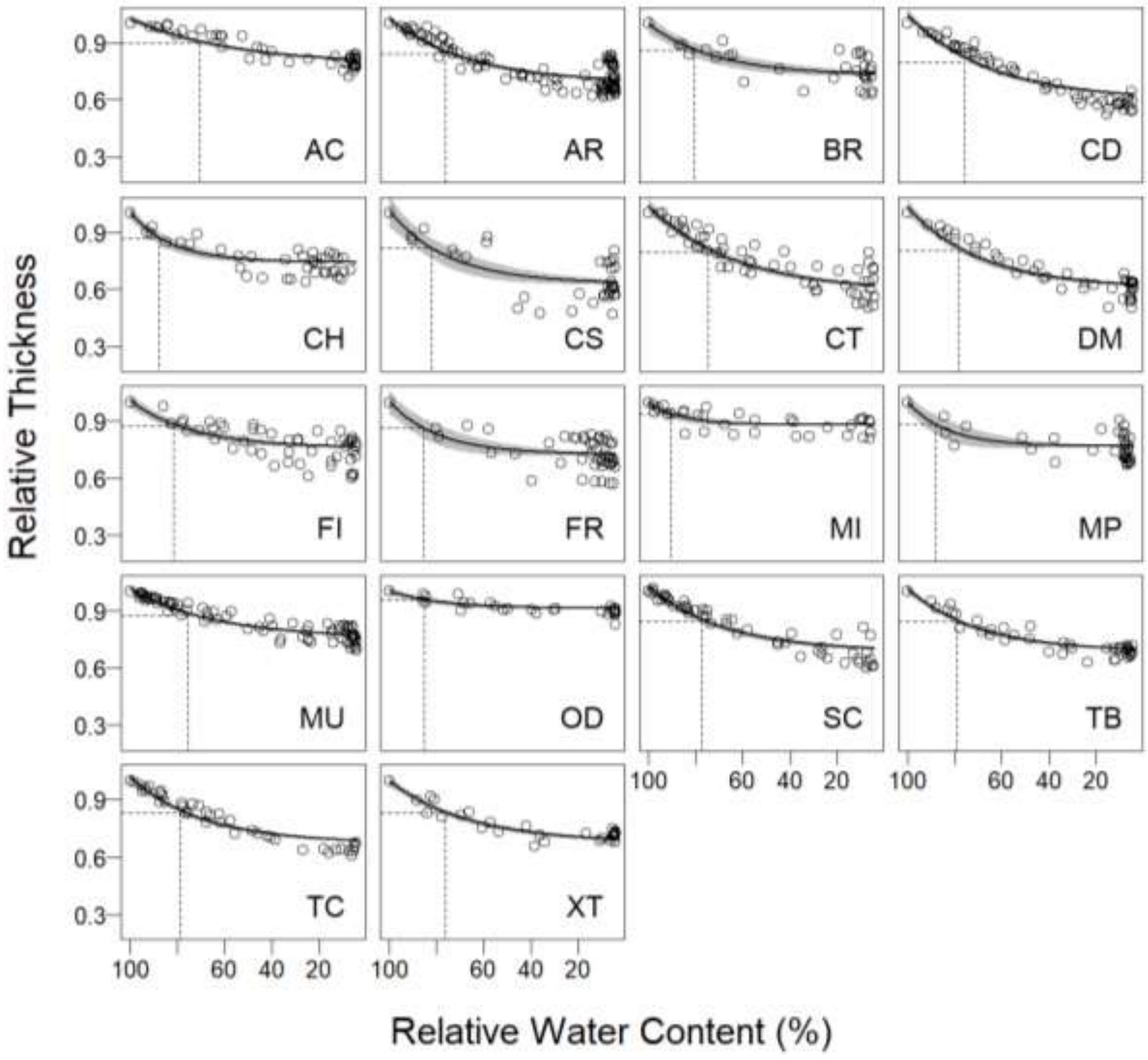


**Figure S4:** Dehydration response curves of the leaf relative thickness for the 18 species studied. Water status of the leaf in terms of leaf relative water content (100 − RWC) ranges from 0% (well hydrated) to 100% (completely dry). Each graph is data pooled from at least 5 replicate individuals. A two parameter exponential decay function fit to the data is shown with grey confidence interval band. The line corresponds to the leaf water status at which the species lost 50% of its shrinkable thickness. The abbreviations for the species are as in table S1.

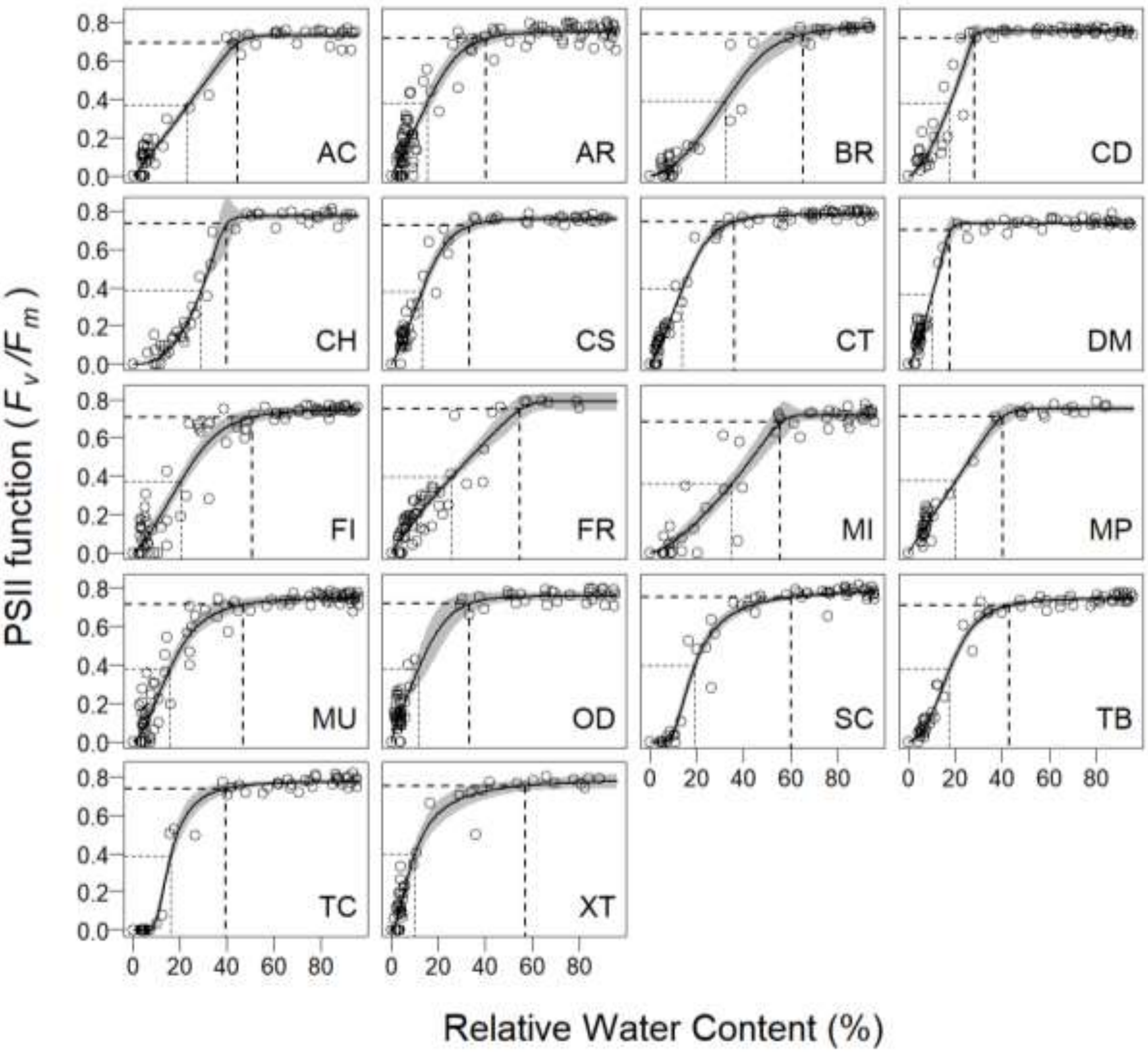


**Figure S5:** Dehydration response curves for leaf PSII function ($F_v/F_m$) that corresponds to dark-adapted chlorophyll A fluorescence efficiency for the 18 species studied. Water status of the leaf in terms of the relative water content (RWC) ranges from 100% (well hydrated) to 0% (completely dry). Each graph is data pooled from at least 5 replicate individuals. A five parameter log-logistic function fit to the data is shown with grey confidence interval band. The lines correspond to the RWC at which 5% loss of PSII function occurs (dashed) and the RWC at which 50% loss of PSII function occurs (solid). The abbreviations for the species are as in table S1.

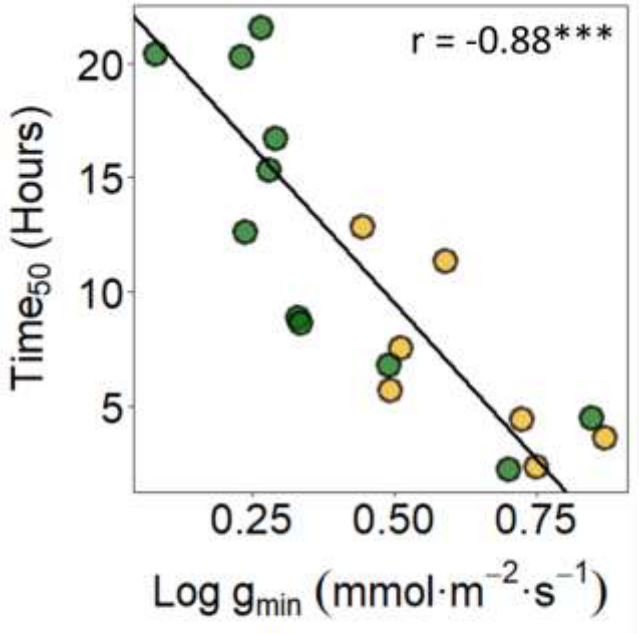


**Figure S6:** Relation between minimum conductance and the time taken to reach 50% leaf RWC for the 18 species studied. Each point corresponds to a species (yellow – deciduous, green – evergreen). A type-2 regression is shown for a significant relationships with Spearman's correlation coefficient (r) corresponding to a *P* < 0.01.

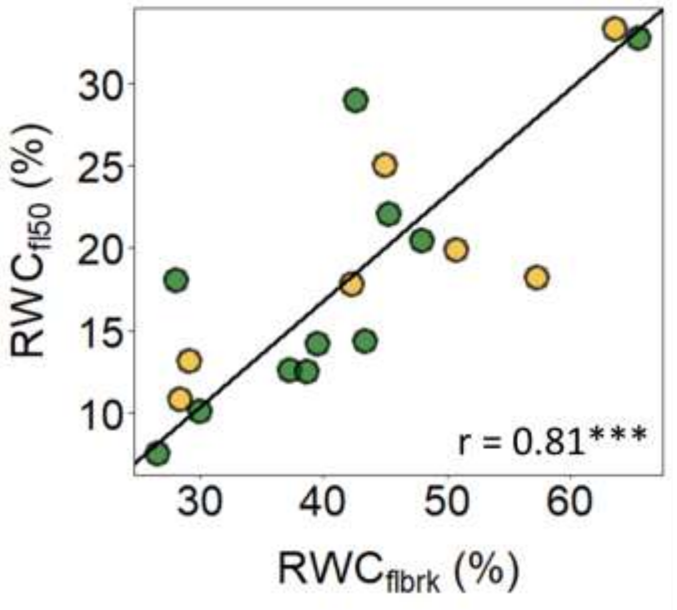


**Figure S7:** Relation between RWC at 50 % decrease in $F_v/F_m$ and RWC at 5 % decrease in $F_v/F_m$ for the 18 species studied. Each point corresponds to a species (yellow – deciduous, green – evergreen). A type-2 regression is shown for the significant relationship with Spearman's correlation coefficient (r) corresponding to a $P < 0.01$.

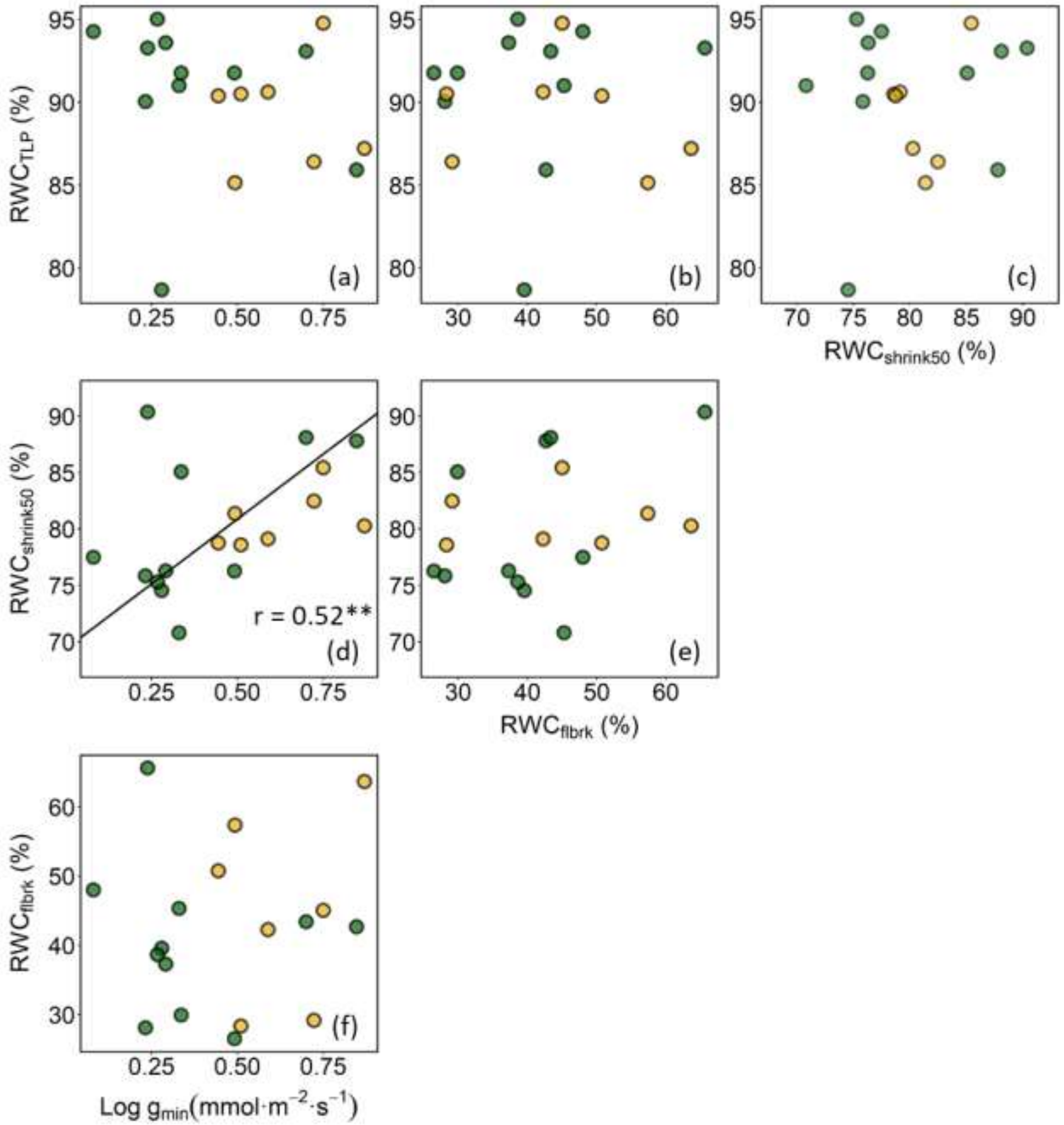


**Figure S8:** Relation between minimum cuticular conductance ($g_{min}$) and the three critical thresholds $RWC_{TLP}$, $RWC_{shrink50}$ and $RWC_{flbrk}$ for the 18 study species. Each point corresponds to a species (yellow – deciduous, green – evergreen). A type-2 regression is shown for the significant relationship with Spearman's correlation coefficient (r) corresponding to a $P < 0.05$.

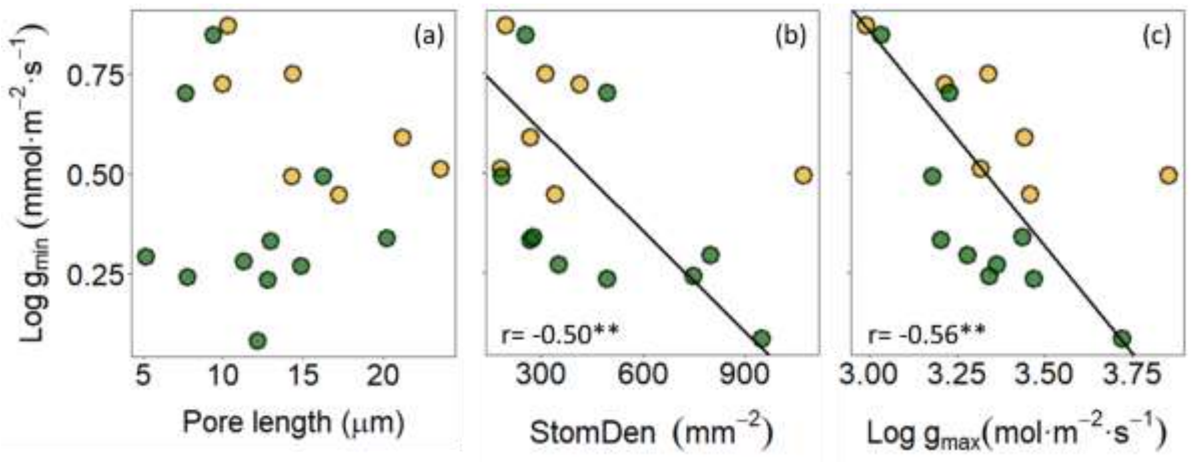


**Figure S9:** Relation between minimum conductance and (a) stomatal pore length, (b) stomatal density and (c) maximum stomatal conductance. Stomatal density and $g_{wmax}$ data could not be estimated for *Callicarpa tomentosa*. Each point corresponds to a species (yellow – deciduous, green – evergreen). A type-2 regression is shown for significant relationships. Solid line corresponds to a relationship with Spearman's correlation coefficient (r) corresponding to a $P < 0.05$.

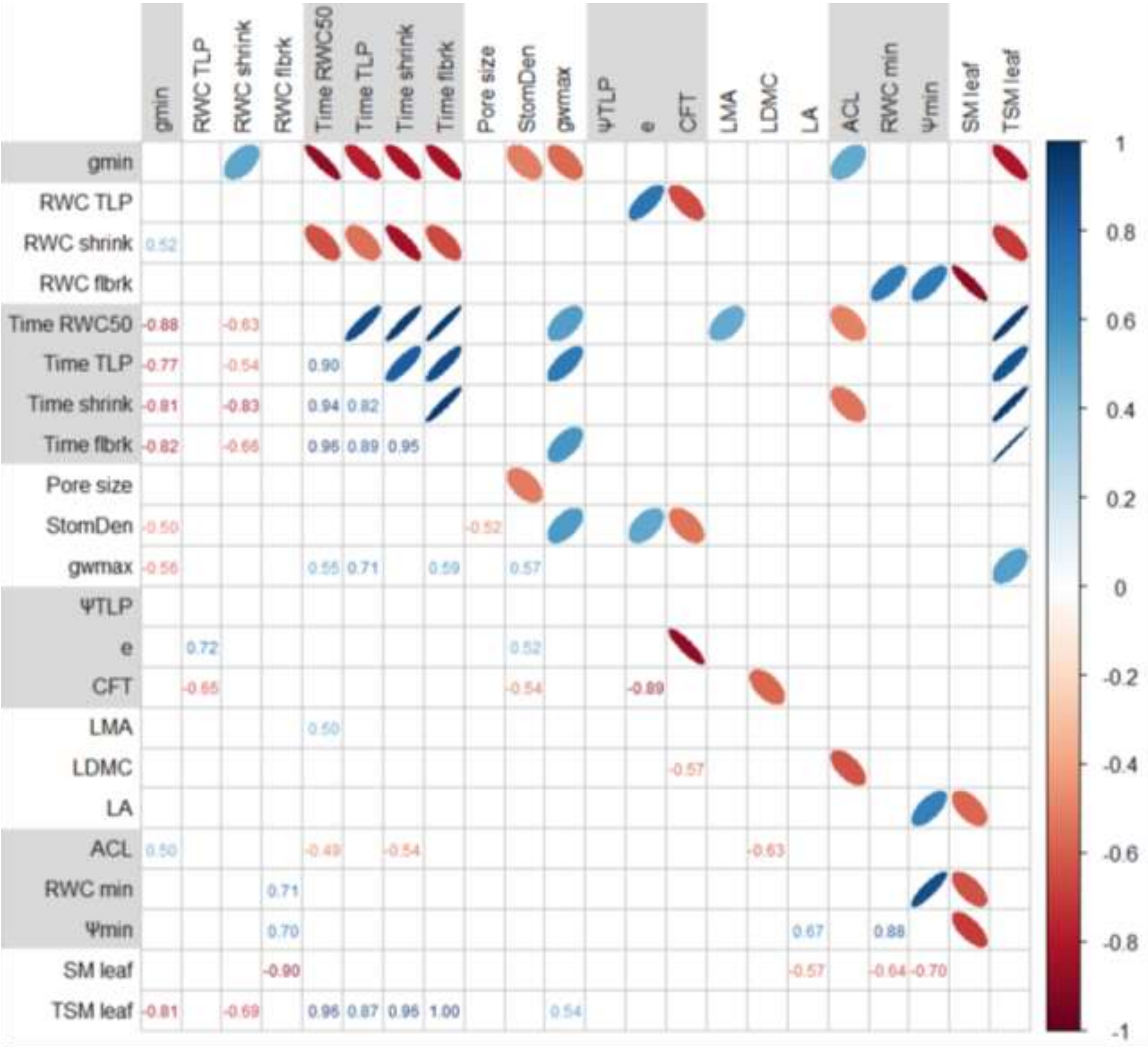


**Figure S10:** Correlation matrix for the hydraulic and functional traits examined in the 18 study species. Spearman's correlation coefficients are presented, and also indicated visually by the eccentricity of the ellipse. Significant relationships with $P < 0.05$ are shown. $SM_{Leaf}$ is the physiological threshold-based safety margin calculated as the difference between the $RWC_{TLP}$ and $RWC_{flbrk}$. $TSM_{leaf}$ is the time-based safety margin calculated as difference between $Time_{TLP}$ and $Time_{flbrk}$. Abbreviations for all other traits are described in Table 1.

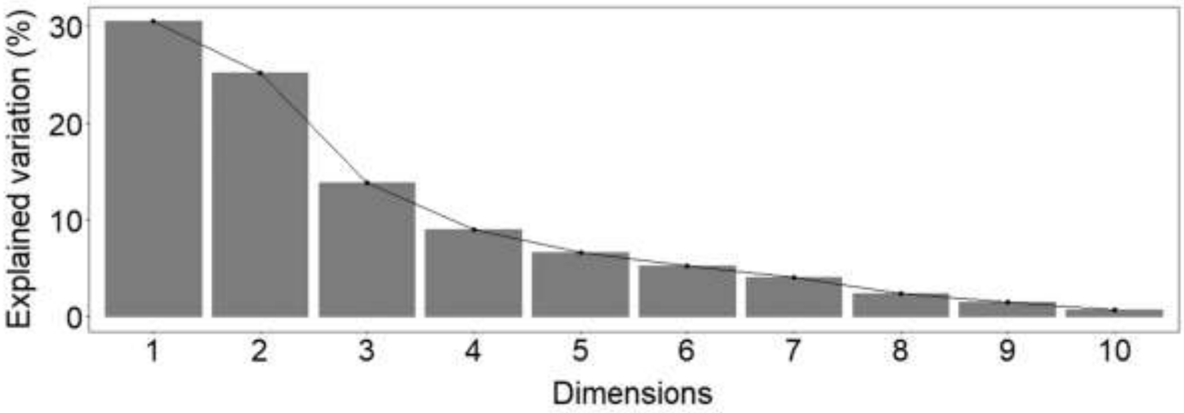


**Figure S11:** Scree plot of the principal component analysis showing the explained variation of each of the dimensions for the 15 leaf traits of the 17 species in this study. *Ficus racemosa* was omitted from this analysis as data for $RWC_{min}$ was not available for this species. Stomatal traits were excluded from the analysis as data was not available for all species. The first three dimensions explained 69.54% of the total variation.

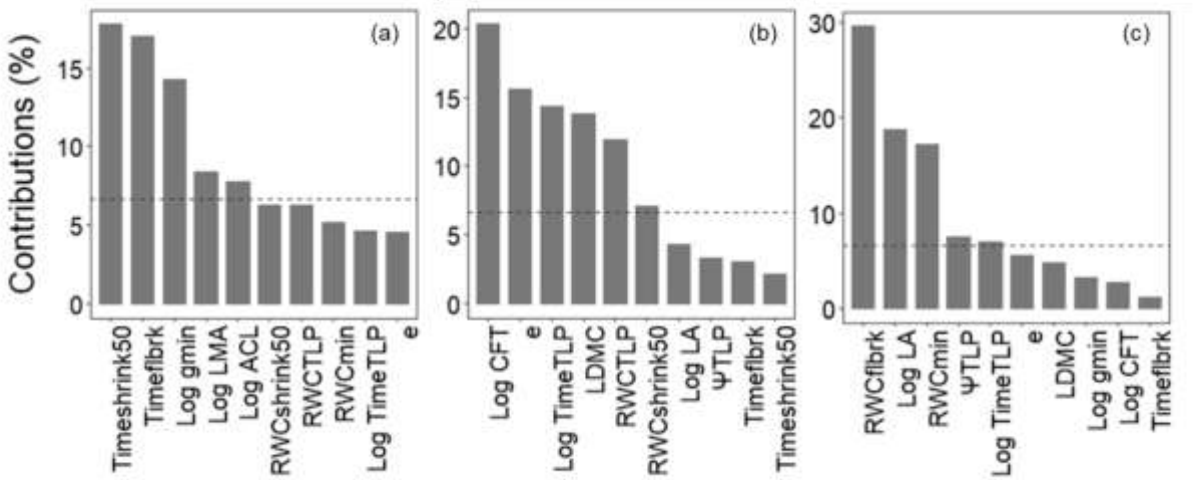


**Figure S12:** Contribution of variables to each of the first three principal component axes; (a) dimension 1, (b) dimension 2 and (c) dimension 3. The dotted line is a reference value which represents the expected values if the contributions of the variables were uniform. Variables above the reference line is considered as important in contributing to the dimension. Abbreviations as in Table 1.

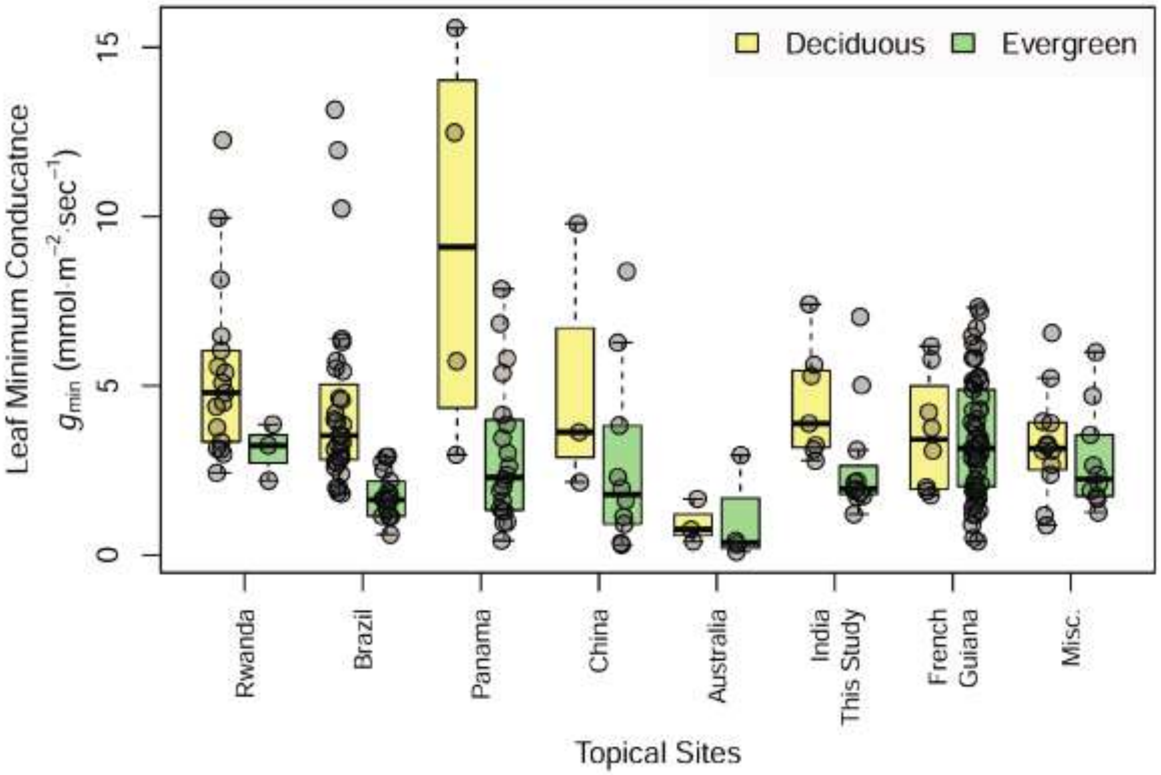


**Figure S13:** Leaf minimum conductance ($g_{min}$) in woody angiosperms from tropical forests. Data were pooled from studies from different sites within regions. Box plots are shown for deciduous (yellow) and evergreen (green) leaf habits, and symbols represent species. (References from which this data were extracted: Boisseaux et al. 2024, Duursma et al. 2016; Levionnes et al. 2021, Loram-Lorenco et al. 2022, Machado et al. 2021, Manzi et al. 2021, Middleby et al. 2024, Schuster et al. 2016; Slot et al. 2021, Witteman et al. 2022, Ziegler et al. 2024).

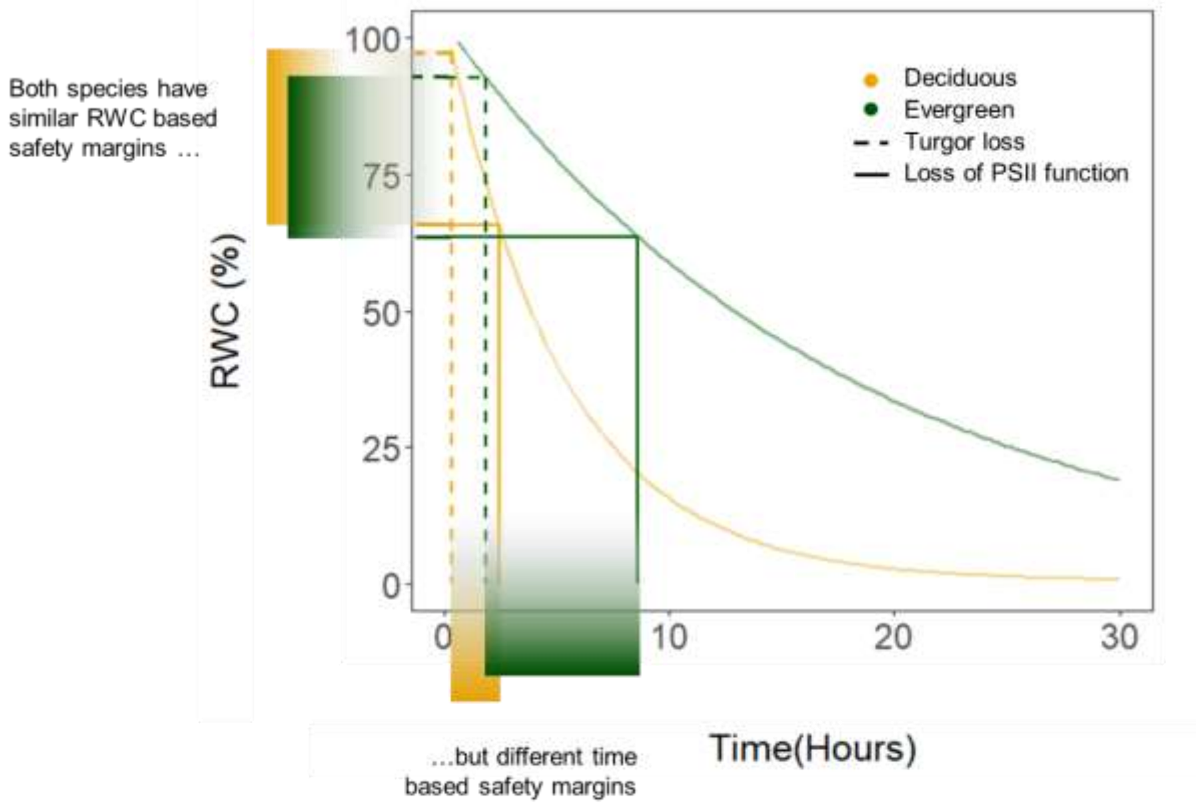


**Figure S14:** Threshold-based and time-based safety margins: Change in leaf RWC with time for leaves of a representative evergreen species (*Mangifera indica*) and a deciduous species (*Bridelia retusa*) during the course of the dehydration assay. Physiological threshold-based safety margin ($SM_{leaf}$) calculated as the difference between $RWC_{tlp}$ and $RWC_{flbrk}$, and time-based safety margin ($TSM_{leaf}$), calculated as the difference between time to $RWC_{tlp}$ ($Time_{TLP}$) and time to $RWC_{flbrk}$ ($Time_{flbrk}$), are shown on the y-axis and x-axis, respectively. Though the physiological threshold-based safety margins are similar for the two species, the time-based safety margins is bigger for the evergreen species, primarily due to its lower dehydration rate (lower $g_{min}$).
